# scSTEC: Annotating Chromatin Subcompartments from Individual Single-Cell Hi-C Data via Direct Two-Dimensional Embedding and Chromosome-wise Consensus

**DOI:** 10.64898/2026.10.28.762654

**Authors:** Yongli Peng, Liyuan Yang, Hao Ge

## Abstract

**Motivation:** Single-cell Hi-C (scHi-C) technologies enable the study of three-dimensional (3D) genome organization at individual-cell resolution. Many computational methods have been developed to infer hierarchical structures from scHi-C data. However, annotating chromatin subcompartments from sparse scHi-C data remains a challenge. The recently developed scGHOST provides a framework for single-cell subcompartment annotation, but relies on a deep learning-based architecture, a multi-step upstream pipeline involving cell embedding and matrix imputation, and cross-cell information sharing. These features complicate algorithmic interpretation and make the annotation of an individual cell dependent on information from other cells in the dataset. Hence, methods are needed that annotate subcompartments directly from an individual cell’s own contact matrix, without complex upstream processing or cross-cell information sharing.

**Results:** We developed scSTEC, a computational framework that annotates subcompartments directly from an individual scHi-C matrix without data imputation, complex upstream processing, or cross-cell data sharing. We show that the estimated inter-chromosomal matrix constructed by scGHOST is intrinsically of rank 2, revealing that the geometry relevant to clustering can be represented in two dimensions. Leveraging this low-dimensional structure, scSTEC performs direct subcompartment annotation through 2D genomic embedding and chromosome-wise consensus. Across bulk Hi-C, downsampled simulations, and real scHi-C datasets (four human cell lines, adult mouse cortex, developing mouse embryos, and developing olfactory sensory neurons), scSTEC achieves subcompartment annotations comparable to scGHOST despite operating on each cell independently, while providing greater algorithmic interpretability. Furthermore, subcompartments identified by scSTEC effectively resolve cell types, capture developmental dynamics, and show consistent associations with gene expression and epigenetic profiles.

**Availability:** scSTEC is freely available at https://github.com/Ge-lab-pku/scSTEC.

## 1 Introduction

The three-dimensional (3D) organization of the genome within the nucleus plays a fundamental role in gene regulation, DNA replication, and cellular function [1, 2, 3]. Chromosome conformation capture techniques [4], particularly Hi-C [1], enable genome-wide mapping of chromatin interactions, revealing a hierarchical organization that includes chromosome territories [5], A/B compartments [1], topologically associating domains (TADs) [6, 7], and chromatin loops [8]. At the megabase scale, the genome partitions into two major compartments: the active, gene-rich A compartment, associated with euchromatin; and the inactive, gene-poor B compartment, associated with heterochromatin [1]. Beyond these binary A/B compartments, higher-resolution Hi-C analyses have further refined this structure by identifying chromatin subcompartments. Rao et al. [8] clustered on the inter-chromosomal Hi-C matrix to divide A/B compartments into five subcompartments (A1, A2, B1, B2, B3), each characterized by unique long-range contact patterns and distinct epigenetic signatures, such as specific histone modifications and replication timing. These subcompartments provide a more nuanced view of the 3D genome’s functional architecture [8].

The recent emergence of single-cell Hi-C (scHi-C) technologies enables the study of 3D genome organization at individual-cell resolution [9, 10, 11, 12], opening new avenues for investigating cell-to-cell heterogeneity in chromatin organization. However, scHi-C data are inherently sparse and noisy, posing substantial challenges for reliably detecting chromatin structures in single cells [9, 13, 14, 15, 16]. While several computational methods can annotate A/B compartments from scHi-C matrices [12, 17, 18], annotating subcompartments at the single-cell level remains challenging due to the difficulty of capturing reliable inter-chromosomal contacts. Recently, scGHOST [19] was developed to address this by leveraging Higashi-based imputation to estimate inter-chromosomal matrices, followed by GHMM or K-means clustering. While scGHOST represents an important advance, its reliance on deep neural network training and a multi-step upstream pipeline complicates algorithmic interpretation. Moreover, its use of cross-cell information means that subcompartment annotation for an individual cell is not based solely on that cell’s own Hi-C matrix.

Here, we introduce scSTEC (Single-cell Subcompartment Annotation via Two-dimensional Embedding and Chromosome-wise Consensus), a computational framework for direct subcompartment annotation from individual scHi-C matrices. scSTEC analyzes each cell independently: subcompartment annotation for a given cell relies on its own Hi-C matrix, requiring no imputation, complex upstream analyses (such as cellular embeddings or A/B compartment inference), or cross-cell data sharing. After valid genomic bins are identified, the core annotation procedure operates independently on each cell, rather than deriving cell-specific annotations through joint analysis of a collection of cells.

The foundation of scSTEC rests on the observation that the construction of the estimated inter-chromosomal matrix used by scGHOST is intrinsically of rank 2. This low-rank structure reveals that the geometry underlying the clustering of genomic bins in this high-dimensional representation can be captured in two dimensions, thereby avoiding the need to explicitly construct and cluster the high-dimensional matrix. Specifically, scSTEC first represents each genomic locus by a two-dimensional vector, whose two coordinates summarize its contact-weighted CpG densities with CpG-defined A-like and B-like genomic bins, respectively. This representation is then transformed into a 2D embedding using individual chromosomes as references. A chromosome-wise consensus mechanism then integrates the resulting annotations across reference chromosomes to obtain the final subcompartment assignments and corresponding consensus scores.

We validated scSTEC on bulk Hi-C [8], simulated scHi-C, and various real scHi-C datasets, including four human cell lines [20, 21], adult mouse cortex [22, 23], developing mouse embryos [22], and developing olfactory sensory neurons [20]. Despite operating on each cell independently and without complex upstream feature extraction, scSTEC achieved subcompartment annotations comparable to scGHOST and remained robust under severe data sparsity. Furthermore, scSTEC identified cell-type-specific subcompartment patterns, captured developmental transitions, and revealed consistent associations between subcompartment organization, gene expression, and epigenetic profiles.

## 2 Methods

The overall architecture of scSTEC is summarized in Figure 1. Designed to annotate subcompartments at single-cell resolution, scSTEC operated directly on intra-chromosomal contact matrices of query and reference chromosomes within the same nucleus (Figure 1). Genomic regions were partitioned into non-overlapping bins at a chosen resolution (e.g., 500 kb). The algorithm accepted both bulk Hi-C and scHi-C matrices as inputs.

**Figure 1:**
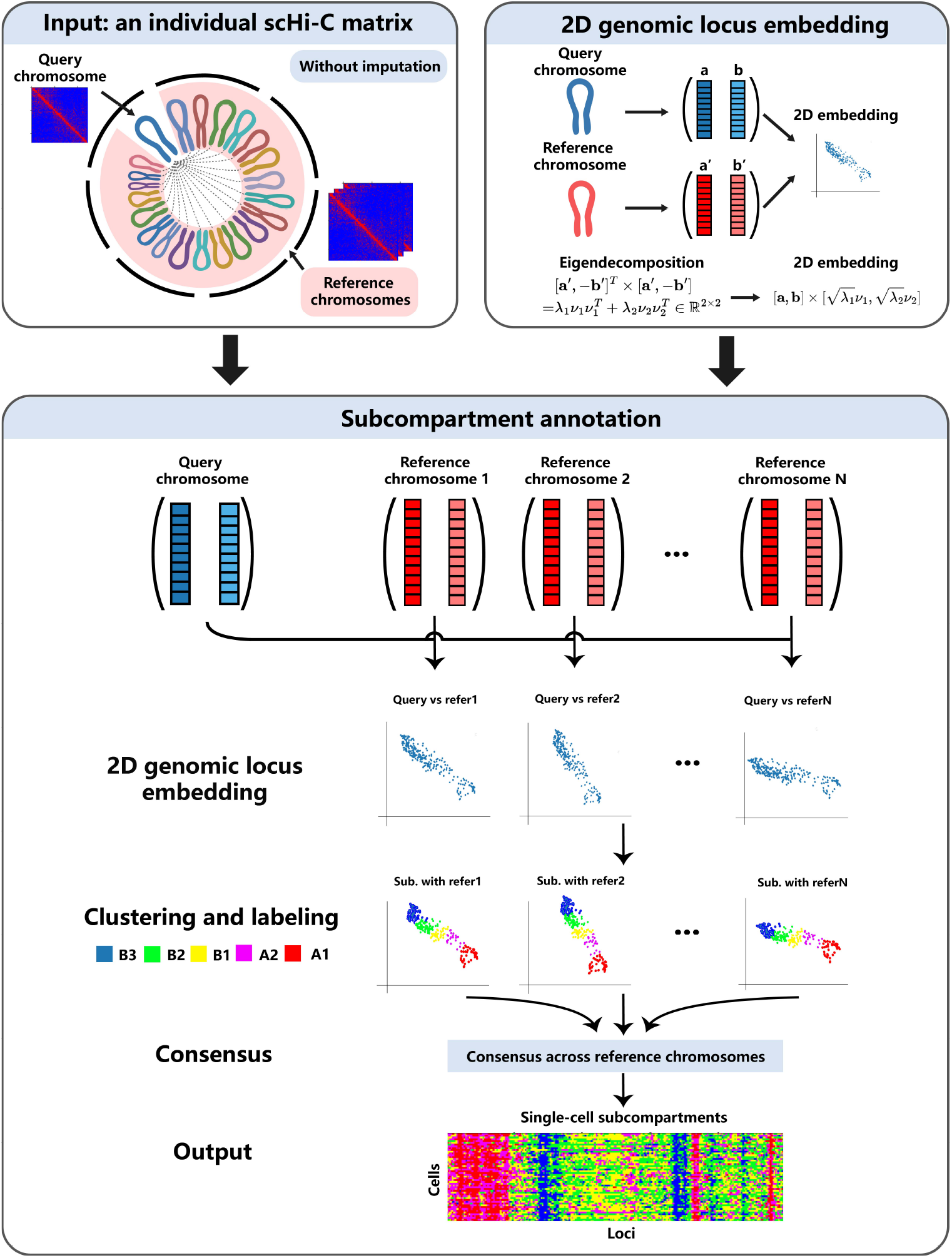
Overview of the scSTEC framework. scSTEC takes the intra-chromosomal scHi-C contact matrices of query and reference chromosomes from the same cell as input. scSTEC projects each query chromosome into a 2D embedding using a transformation defined by a reference chromosome. Clustering and subcompartment labeling are then performed in this 2D space. Finally, annotations obtained from different reference chromosomes are integrated through chromosome-wise consensus to produce the final subcompartment annotation.

We first computed a 2D genomic representation, 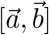, whose two coordinates summarized each locus’s contact-weighted CpG densities with CpG-defined A-like and B-like genomic bins, respectively. The two-dimensional formulation was motivated by our mathematical analysis of the construction of the estimated inter-chromosomal matrix used by scGHOST (Figure 1). We then transformed this representation into a 2D genomic embedding through an eigenvector-guided projection defined by each reference chromosome. Next, we performed clustering and labeling on these 2D embeddings. Finally, we established a consensus labeling across all reference chromosomes to obtain the final subcompartment annotation of each genomic bin.

In the following sections, we introduced the scSTEC workflow for scHi-C data. The processing steps for bulk Hi-C were similar and are detailed in the Supplementary Material.

### 2.1 Two-dimensional genomic representation and embedding

Inspired by scGHOST and scA/B values, scSTEC first computed a 2D representation, 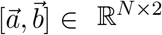, for a query chromosome with *N* genomic bins, where (*a*_*i*_, *b*_*i*_) represented locus *i*:

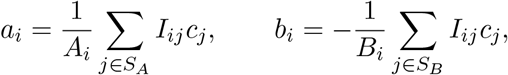

where *I* was the raw scHi-C matrix, *c* was the CpG density, and

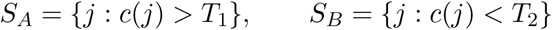

denoted the sets of high- and low-CpG bins, respectively. We used the top and bottom 30% of bins ranked by CpG density to define *S*_*A*_ and *S*_*B*_. Here, 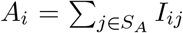 and 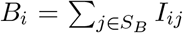. The CpG density *c* was normalized to [ −0.5, 0.5]. Because *c*_*j*_ *>* 0 for *j* ∈ *S*_*A*_ and *c*_*j*_ *<* 0 for *j* ∈ *S*_*B*_, the negative sign in the definition of *b*_*i*_ placed the two coordinates on the same positive orientation. Intuitively, *a*_*i*_ and *b*_*i*_ summarized the contact-weighted CpG densities of locus *i* with high- and low-CpG genomic bins, respectively. A larger *a*_*i*_ indicated a stronger tendency of locus *i* toward the A compartment, whereas a larger *b*_*i*_ indicated a stronger tendency toward the B compartment.

Direct clustering of 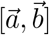 performed worse than the scGHOST framework on simulated scHi-C datasets (Supplementary Figure S1). We therefore constructed a 2D genomic embedding that preserves the relevant geometry of the estimated inter-chromosomal matrix used by scGHOST. In the scGHOST framework, for a query chromosome with *N* bins and a reference chromosome with *N* ^*′*^ bins, whose 2D representations were 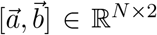 and 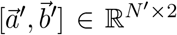, respectively, the estimated inter-chromosomal matrix is constructed as

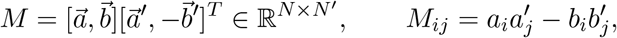

where *i* = 1, …, *N* and *j* = 1, …, *N* ^*′*^. scGHOST then applies a GHMM (Gaussian hidden Markov model) or K-means clustering to the rows and columns of *M* for subcompartment annotation.

Because 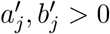, for a fixed *j*,

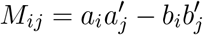

was linear in *a*_*i*_ and *b*_*i*_, increasing with *a*_*i*_ and decreasing with *b*_*i*_. Typically, *a*_*i*_ and *b*_*i*_ were negatively correlated. Thus, larger values of *M*_*ij*_ tended to correspond to a more A-like state of the row locus *i*. The same reasoning applied to the reference locus *j*. This construction was therefore expected to distinguish *A*–*A, A*–*B*, and *B*–*B* interactions, with *A*–*A* pairs tending to have larger *M*_*ij*_ values and *B*–*B* pairs smaller values.

Mathematically, however, *M* was a rank-2 matrix. Thus, although each row of *M* was represented as an *N* ^*′*^-dimensional vector, its geometry was intrinsically two-dimensional. Explicit clustering in this high-dimensional representation was therefore unnecessary. As shown below, the Euclidean distances between rows of *M* could be represented exactly in a two-dimensional space.

We then considered the squared Euclidean distance between rows *i* and *j* of *M*:

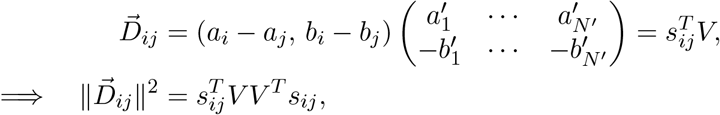

where

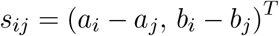

and

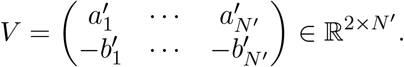

In contrast, direct clustering on 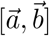 used the ordinary Euclidean distance,

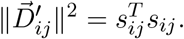

Since *V V* ^*T*^ ∈ ℝ^2*×*2^, let (*λ*_1_, *λ*_2_) and (*ν*_1_, *ν*_2_) denote its eigenvalues and corresponding eigen-vectors, respectively. The distance in scGHOST’s framework could then be rewritten exactly as

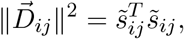

where

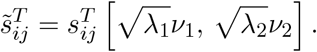

Thus, the Euclidean geometry of the *N* ^*′*^-dimensional rows of *M* could be represented exactly in a two-dimensional space.

Thus, clustering on 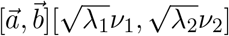 was mathematically equivalent, in terms of Euclidean geometry, to clustering the rows of *M* in the scGHOST framework. Accordingly, in scSTEC, we defined

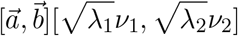

as the 2D genomic locus embedding and performed clustering directly in this two-dimensional space. When *λ*_1_ = *λ*_2_, this transformation was a scaled isometry, and the clustering results remained invariant. In practice, however, the ratio *λ*_1_*/λ*_2_ was typically large (Supplementary Figure S2) and varied across reference chromosomes, whereas the eigenvectors *ν*_1_ and *ν*_2_ were highly consistent across chromosomes. As shown in Supplementary Figure S2, the cosine similarity between *ν*_1_ of each chromosome and that of chromosome 1 exceeded 0.98 in most cases. These results indicated that differences in the 2D transformations induced by different reference chromosomes were primarily driven by the eigenvalues *λ*_1_ and *λ*_2_.

### 2.2 Consensus across chromosomes

After constructing the 2D embeddings, we performed clustering of genomic bins and labeled the five clusters based on CpG density. We then established a consensus across reference chromosomes to obtain the final subcompartment assignments.

For a query chromosome *n*, we computed the 2D embedding

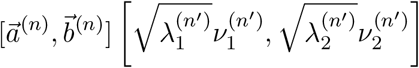

for each reference chromosome *n*^*′*^ ≠ *n*. We applied K-means clustering (*k* = 5) to each 2D embedding and labeled the five clusters as A1, A2, B1, B2, and B3 according to their decreasing mean CpG densities.

Because the 2D embedding of a query chromosome depended on the choice of reference chromosome, different reference chromosomes could yield slightly different clustering and labeling results. We therefore introduced a chromosome-wise consensus procedure to obtain a robust final annotation. For each query chromosome *n*, the clustering and labeling procedure was repeated *N*_*c*_ − 1 times, once for each reference chromosome *n*^*′*^ ≠ *n*, where *N*_*c*_ was the total number of chromosomes (22 for human and 19 for mouse). For each locus, we determined its final subcompartment assignment as the class most frequently assigned across all reference chromosomes. We also defined a consensus score as the proportion of reference chromosomes yielding the final assignment, which quantified the consistency of the annotation across reference chromosomes. In bulk Hi-C evaluations (Supplementary Figure S3), all bins achieved consensus scores above 50%. A1, B2, and B3 consistently exhibited higher consensus scores than A2 and B1.

### 2.3 Quality control of the data

Before cell-specific quality control, we first removed poorly covered genomic bins at the dataset level. For each dataset at a given resolution, we calculated the total number of contacts involving each bin across all cells and excluded bins with fewer than 10 contacts from subsequent analyses. This step was used only to define a common set of valid genomic bins and did not otherwise pool contact information across cells for subcompartment annotation. Further details are provided in the Supplementary Material.

During the 2D embedding of the GM12878 scHi-C data, we observed that some matrices contained bins with *a*_*i*_ = 0 or *b*_*i*_ = 0. These bins behaved as outliers in the 2D embedding space and distorted the clustering results (Supplementary Figure S4). We therefore excluded these bins during clustering. After clustering, the excluded bins were assigned to the nearest cluster centroid based on their Euclidean distances in the 2D embedding space.

For each cell, if fewer than 20% of genomic bins remained after excluding those with *a*_*i*_ = 0 or *b*_*i*_ = 0, we excluded the entire cell from downstream analysis. This empirical threshold rarely excluded cells in recent high-coverage datasets and had little effect on the downstream analyses. Nevertheless, reliable subcompartment annotation requires a minimum number of informative contacts. Simulated matrices downsampled to 1/6000 retained contact numbers comparable to those in recent datasets such as LiMCA (Supplementary Figure S5). Based on the LiMCA data at 500 kb resolution, we used an empirical QC guideline that excluded cells with matrix sparsity above 0.98 or fewer than 3,000 total contacts (Supplementary Figure S5). These thresholds were empirical and should be adjusted according to dataset coverage and resolution.

### 2.4 Simulated scHi-C matrices

We generated simulated scHi-C datasets by downsampling the bulk GM12878 Hi-C matrix at 500 kb resolution, using the subcompartment annotations obtained with the original method of Rao et al. [8] as the ground truth. For a given downsampling rate 1*/r*, we binomially downsampled each contact count in the bulk Hi-C matrix with sampling probability 1*/r* to generate a simulated scHi-C contact map. We repeated the downsampling 50 times at each rate. The downsampling rates were 1/10, 1/100, 1/1000, 1/2500, 1/4000, and 1/6000.

### 2.5 Real scHi-C and scRNA-seq datasets

We analyzed publicly available scHi-C datasets covering human cell lines, adult mouse brain, mouse embryonic development, and developing olfactory sensory neurons. The datasets were processed largely following the pipelines of the original studies, with additional procedures required for subcompartment annotation and downstream analyses. For bulk and pseudo-bulk analyses, contact maps were processed using `cooler` and `cooltools`, and A/B compartments were identified using `cooltools`. scRNA-seq data were preprocessed and clustered using Seurat, with marker genes identified using FindAllMarkers. Dataset-specific preprocessing procedures and parameter settings are described in the Supplementary Material.

### 2.6 Comparing scSTEC with other methods

We compared scSTEC with the GHMM-based method of Rao et al.(hereafter, Rao’s method) and scGHOST[8, 19]. Because the GHMM-based method was developed for bulk Hi-C data, we applied it only to the bulk and simulated Hi-C matrices. For scGHOST, we used Higashi for matrix imputation and A/B compartment inference and then ran the full scGHOST pipeline with default parameters. Agreement between subcompartment annotations was quantified using Spearman correlations based on subcompartment labels and the intersection score defined as the fraction of loci assigned to the same subcompartment by the two methods.

### 2.7 Pooled single-cell subcompartment annotations

We obtained pooled single-cell subcompartment annotations by aggregating independently inferred subcompartment labels across cells. Specifically, for each genomic bin, we assigned the subcompartment label most frequently observed across cells:

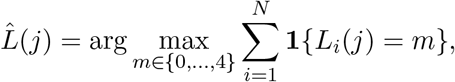

where *L*_*i*_(*j*) denotes the subcompartment label of genomic bin *j* in cell *i, N* is the number of cells included in the pooling, and the five subcompartments B3, B2, B1, A2, and A1 are encoded as 0, 1, 2, 3, and 4, respectively.

### 2.8 Stable and variable subcompartments

For each genomic locus, we quantified subcompartment variability across cells using the entropy of the observed subcompartment distribution (see Supplementary Material). Loci in the bottom and top 30% of the variability distribution were classified as stable and variable loci, respectively. We further evaluated the robustness of the downstream results using alternative percentile cutoffs of 5%, 10%, 25%, and 50% (Supplementary Figure S6).

## 3 Results

### 3.1 scSTEC applied to bulk Hi-C and simulated scHi-C matrices

We first evaluated scSTEC on the GM12878 bulk Hi-C matrix [8]. Because scGHOST required multiple samples for model training, we generated 50 samples by independently downsampling the bulk matrix at a rate of 1/10 and used them to train scGHOST. As shown in Figure 2A, both visual comparison and quantitative metrics (intersection and correlation) showed high agreement between scSTEC and scGHOST. This result was consistent with our mathematical derivation showing that the Euclidean geometry of scGHOST’s estimated inter-chromosomal matrix could be represented exactly in two dimensions.

**Figure 2:**
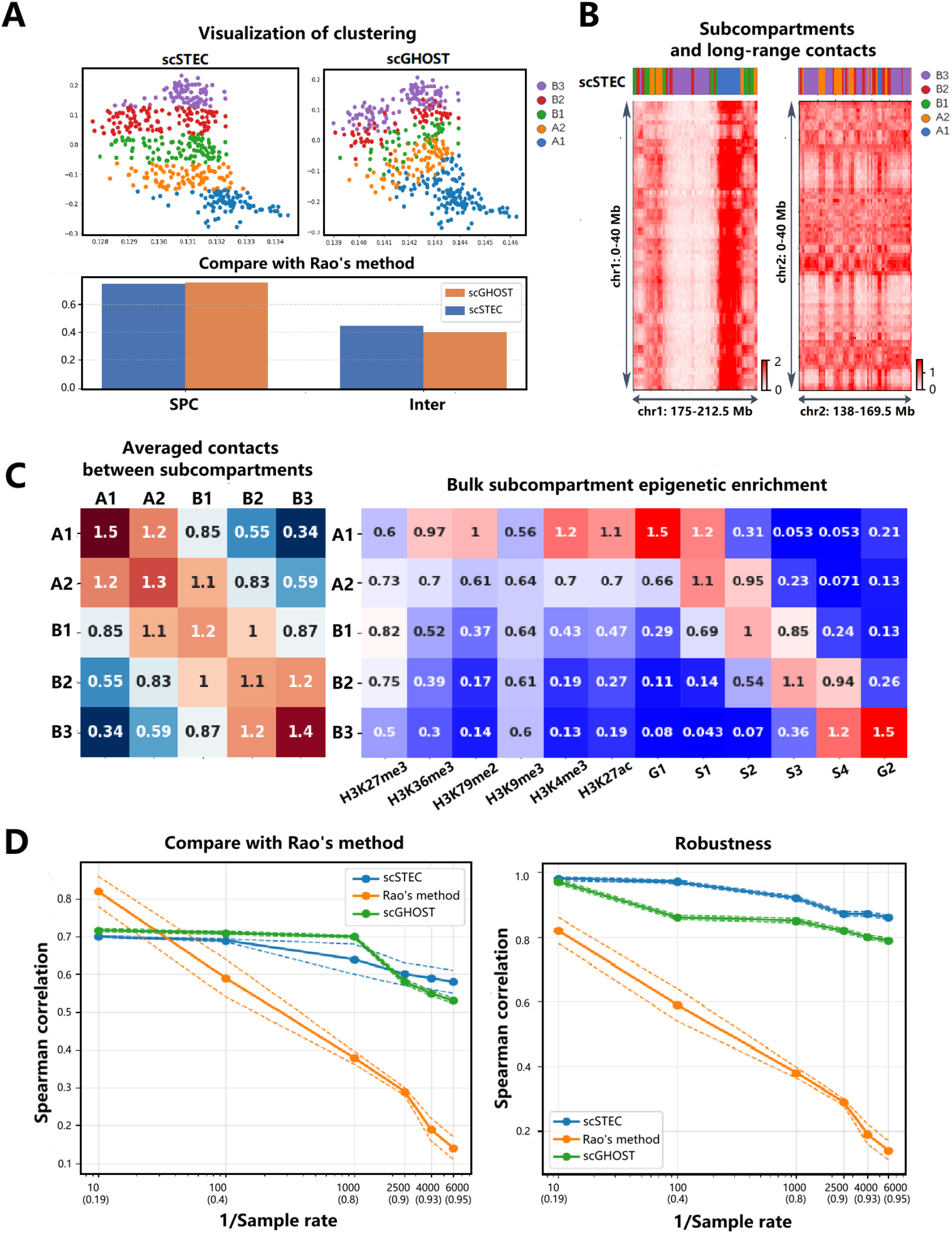
Performance of scSTEC on bulk and simulated Hi-C data. (A) Visualization of the five subcompartment annotations obtained by scSTEC and comparison with Rao’s method using Spearman correlation (SPC) and intersection (Inter). (B) Visualization of scSTEC subcompartment annotations together with long-range contact patterns in the O/E-normalized Hi-C map. (C) Mean O/E-normalized contact frequencies between pairs of subcompartments and median fold changes of epigenomic signals over controls across the five subcompartments. (D) Comparison of scSTEC, Rao’s method, and scGHOST on simulated scHi-C matrices across different downsampling rates. Solid lines indicate the mean across 50 independent downsampling replicates, and dashed lines indicate the mean ± standard error. The corresponding sparsity at each downsampling rate is also shown. Robustness was measured by the Spearman correlation between each method’s annotations from down sampled data and its corresponding annotations from the full-depth data.

In Supplementary Figure S7, we further examined the consistency between the five subcompartments and conventional A/B compartments. For each subcompartment, consistency was defined as the fraction of its loci belonging to the corresponding conventional A/B compartment (A for A1/A2 and B for B1/B2/B3). As expected, A1 and A2 predominantly corresponded to the A compartment, whereas B1, B2, and B3 predominantly corresponded to the B compartment. Both scSTEC and scGHOST showed high overall consistency with the A/B compartment partition, while Rao’s annotation showed somewhat lower consistency (Supplementary Figure S7). Among individual subcompartments, A1 and B3 showed the highest consistency, whereas B1 showed the lowest. This pattern was also consistent with the chromosome-wise consensus analysis in Supplementary Figure S3, in which A1 and B3 exhibited particularly high annotation consistency across different reference chromosomes. Together, these results suggest that A1 and B3 represent particularly robust subcompartment states that are also most consistent with the classical A/B compartment partition.

In Figure 2B, we examined whether scSTEC’s annotations captured characteristic long-range contact patterns. We focused on two off-diagonal regions of the O/E-normalized Hi-C map and observed distinct long-range interaction patterns among different subcompartments, consistent with previous findings [8, 19]. In Figure 2C, we further calculated the mean O/E-normalized contact frequency between each pair of subcompartments. Loci assigned to the same subcompartment preferentially interacted, with intra-subcompartment contacts significantly stronger than inter-subcompartment contacts (one-sided *P <* 5 *×* 10^−142^). Moreover, contact strength generally decreased as the subcompartment states became more dissimilar. Together, these results showed that scSTEC recapitulated the characteristic long-range contact organization reported for canonical chromatin subcompartments.

In Figure 2C, we further examined the epigenomic and replication-timing profiles of the subcompartments identified by scSTEC. We computed the median fold change over the control for each epigenomic signal within each subcompartment. Both A1 and A2 were enriched for active histone marks, including H3K36me3, H3K79me2, H3K27ac, and H3K4me3, with A2 showing greater enrichment of H3K9me3 than A1. Both exhibited early replication timing, with A1 completing replication earlier in S phase and A2 extending into mid-S phase. In contrast, the B subcompartments were depleted of active histone marks. B1 was enriched for H3K27me3, a mark associated with facultative heterochromatin, and replicated mainly during mid-S phase. B2 and B3 showed little enrichment for these histone marks and replicated predominantly late in S phase. These profiles were consistent with the characteristic epigenetic and replication-timing patterns previously reported for the five subcompartments [8, 19].

We next evaluated robustness to data sparsity using simulated scHi-C datasets with sampling rates ranging from 1/10 to 1/6000. As shown in Figure 2D, at the 1/10 sampling rate, both scSTEC and scGHOST showed high concordance with the ground truth (Spearman’s *r* = 0.7), comparable to Rao’s method (*r* = 0.8). As the sampling rate decreased, the performance of Rao’s method deteriorated rapidly. In contrast, both scSTEC and scGHOST remained robust to increasing sparsity, maintaining Spearman correlations around 0.6 even at the 1/6000 sampling rate. Their overall performance was comparable, with scSTEC slightly outperforming scGHOST under the sparsest conditions (1/4000 and 1/6000). These results demonstrated that scSTEC remained robust under severe data sparsity despite performing subcompartment annotation without cross-cell information sharing.

### 3.2 scSTEC applied to human cell lines

We next evaluated scSTEC on real scHi-C data from human cell lines, focusing on the high-coverage LiMCA dataset for GM12878.

In Figure 3A, we computed the mean O/E-normalized contacts between subcompartments using the pseudo-bulk Hi-C matrix and pooled single-cell subcompartment annotations. Consistent with the bulk Hi-C analysis, contacts between loci assigned to the same subcompartment were significantly stronger than those between different subcompartments (*P*-values below numerical precision). Moreover, contact strength generally decreased as the subcompartment states became more dissimilar, revealing a graded spatial organization among the five subcompartments.

**Figure 3:**
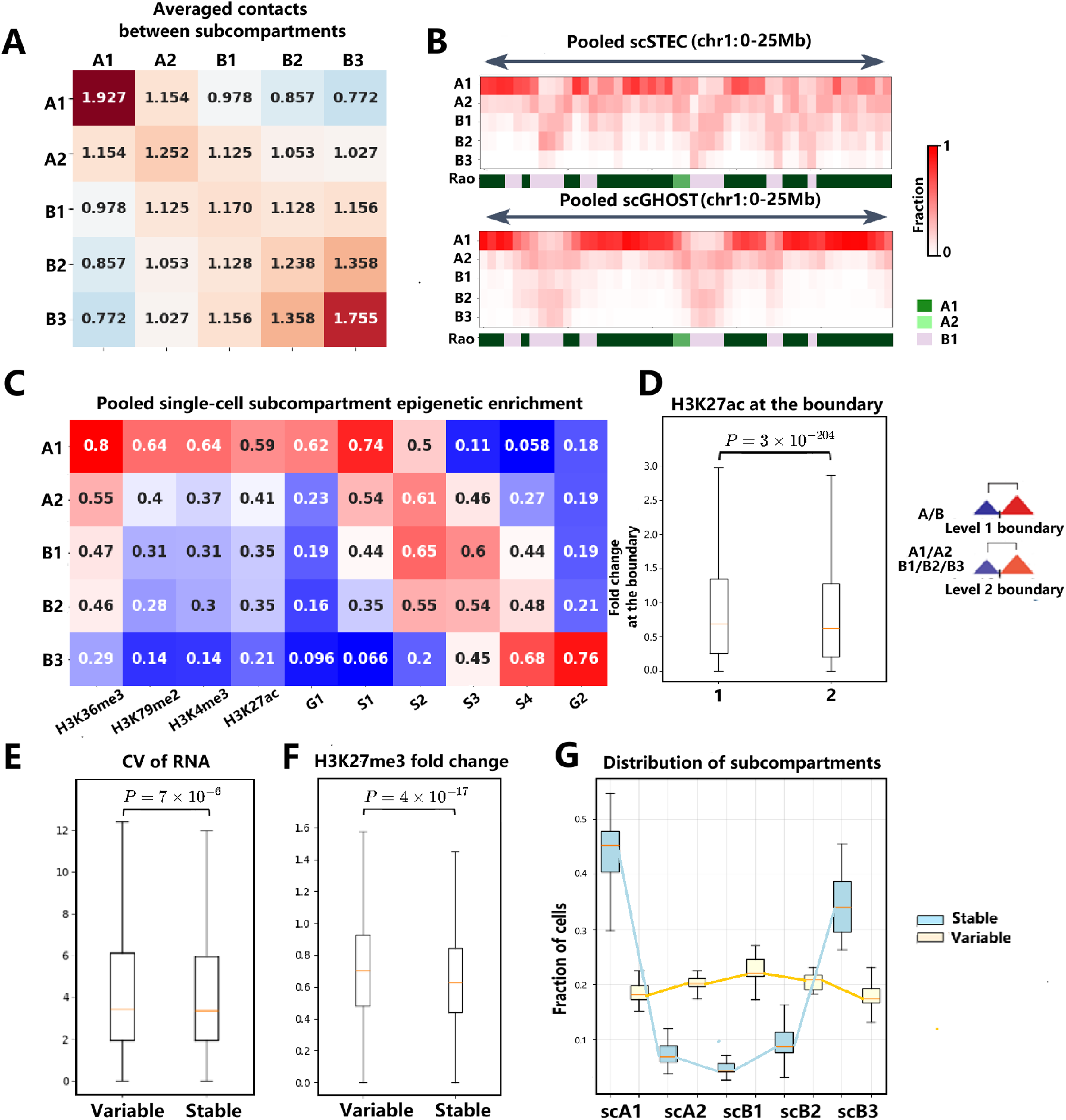
Application of scSTEC to the human GM12878 cell line. (A) Mean O/E-normalized contact frequencies within and between pooled single-cell subcompartments, calculated from the pseudo-bulk Hi-C matrix. (B) Visualization of pooled single-cell subcompartment annotations obtained by scSTEC and scGHOST. (C) Median fold enrichment of epigenomic signals relative to controls within each pooled single-cell subcompartment. (D) Fold changes of H3K27ac across subcompartment boundaries in individual cells. (E) Transcriptional variability at genomic loci with variable and stable subcompartment states in HiRES Ex1 cells. (F) H3K27me3 enrichment at genomic loci with variable and stable subcompartment states in GM12878 cells. (G) Distribution of subcompartment assignments across stable (blue) and variable (yellow) loci genome-wide.

In Figure 3B and Supplementary Figure S9, pooled single-cell subcompartment annotations showed strong agreement with the bulk Hi-C subcompartments reported by Rao et al. [8] and were visually comparable to those obtained by scGHOST. At the genome-wide level, locus pairs assigned to the same pooled scSTEC subcompartment exhibited lower variability in their pseudo-bulk Hi-C contacts than those grouped according to Rao’s method (Supplementary Figure S10). A similar reduction was observed for pooled scGHOST annotations.

In Figure 3C, we examined the epigenomic and replication-timing profiles of the pooled single-cell subcompartments. The results closely recapitulated those observed in the bulk analysis (Figure 2C). Active epigenetic signals were enriched in A1 and A2, and the replication-timing profiles showed similar patterns. We further examined this relationship at the individual-cell level using the four-omics CHARM dataset [23], which simultaneously profiles 3D genome organization and H3K27ac in single cells (Supplementary Figure S11). H3K27ac was enriched in A1, with *P <* 10^−5^ in the vast majority of individual cells, demonstrating that the association between A1 and active chromatin observed at the bulk and pseudo-bulk levels was also detectable within individual cells.

In Figure 3D, we further examined changes in histone marks at subcompartment boundaries. Following Calder [24], we classified boundaries as level 1 (between A and B compartments) or level 2 (within A or B compartments). Histone marks showed larger fold changes across level 1 boundaries than across level 2 boundaries at the individual-cell level, as demonstrated for H3K27ac (Figure 3D), with consistent results for other histone marks and in the pooled analysis (Supplementary Figure S12).

We next characterized cell-to-cell variability in subcompartment organization (Figure 3E-G). Adapting the variability framework developed in our previous work [18], we classified genomic loci as stable or variable using the bottom and top 30% of the subcompartment variability distribution, respectively (Methods). In the HiRES Ex1 dataset, variable loci exhibited significantly greater transcriptional variability than stable loci, as quantified by the coefficient of variation (CV) of RNA expression (Figure 3E, *P* = 7 *×* 10^−6^). We next examined H3K27me3, a histone mark enriched in B1 and associated with facultative heterochromatin [25]. Variable loci showed significantly higher H3K27me3 levels than stable loci across the cell population (Figure 3F, *P* = 4 ∗ 10^−17^). Moreover, H3K27me3 levels were positively correlated with subcompartment variability (Pearson’s *r* = 0.34). This association was consistent with the cell-type-dependent expression patterns previously observed for H3K27me3-repressed genes [26]. Finally, the higher transcriptional variability of variable loci remained consistent across alternative percentile cut-offs (Supplementary Figure S6).

In Figure 3G, we further examined the distribution of single-cell subcompartment assignments for stable and variable loci. Stable loci were predominantly assigned to A1 and B3 (Figure 3G, blue), consistent with the high consensus scores of these subcompartments across reference chromosomes (Supplementary Figure S3). In contrast, variable loci showed a broader distribution across all five subcompartments, with a modest preference for scB1 (Figure 3G, yellow). We further examined the contact patterns of stable and variable loci using the pseudo-bulk Hi-C matrix (Supplementary Figure S13). Stable loci exhibited higher mean O/E-normalized contact frequencies with A1 and B3 subcompartments than variable loci, whereas the opposite pattern was observed for A2, B1, and B2.

The low-dimensional representation used by scSTEC facilitates subcompartment annotation at relatively high resolutions when sufficient contact information is available. In the high-coverage LiMCA and scMicro-C datasets examined here, scSTEC remained practically applicable at resolutions of approximately 50–100 kb, based on empirical QC and cross-resolution consistency criteria (Supplementary Figures S14 and S15).

### 3.3 scSTEC applied to the adult mouse cortex

We next applied scSTEC to scHi-C data from the adult mouse cortex in HiRES dataset [22] to assess whether its subcompartment annotations capture cell-type-specific chromatin organization and are associated with cell-type-specific gene expression. We first generated UMAP embeddings using only the scSTEC subcompartment annotations. The embedding revealed four major cell clusters (Figure 4A): Ex1–3 and In2 cells formed one large cluster, whereas In1, astrocytes, and oligodendrocytes formed three separate clusters. Despite using subcompartment annotations alone and without imputation, scSTEC produced a cellular embedding comparable to those obtained with scHiCluster and scGHOST. These results indicate that scSTEC subcompartment annotations retain substantial cell-type-specific information.

**Figure 4:**
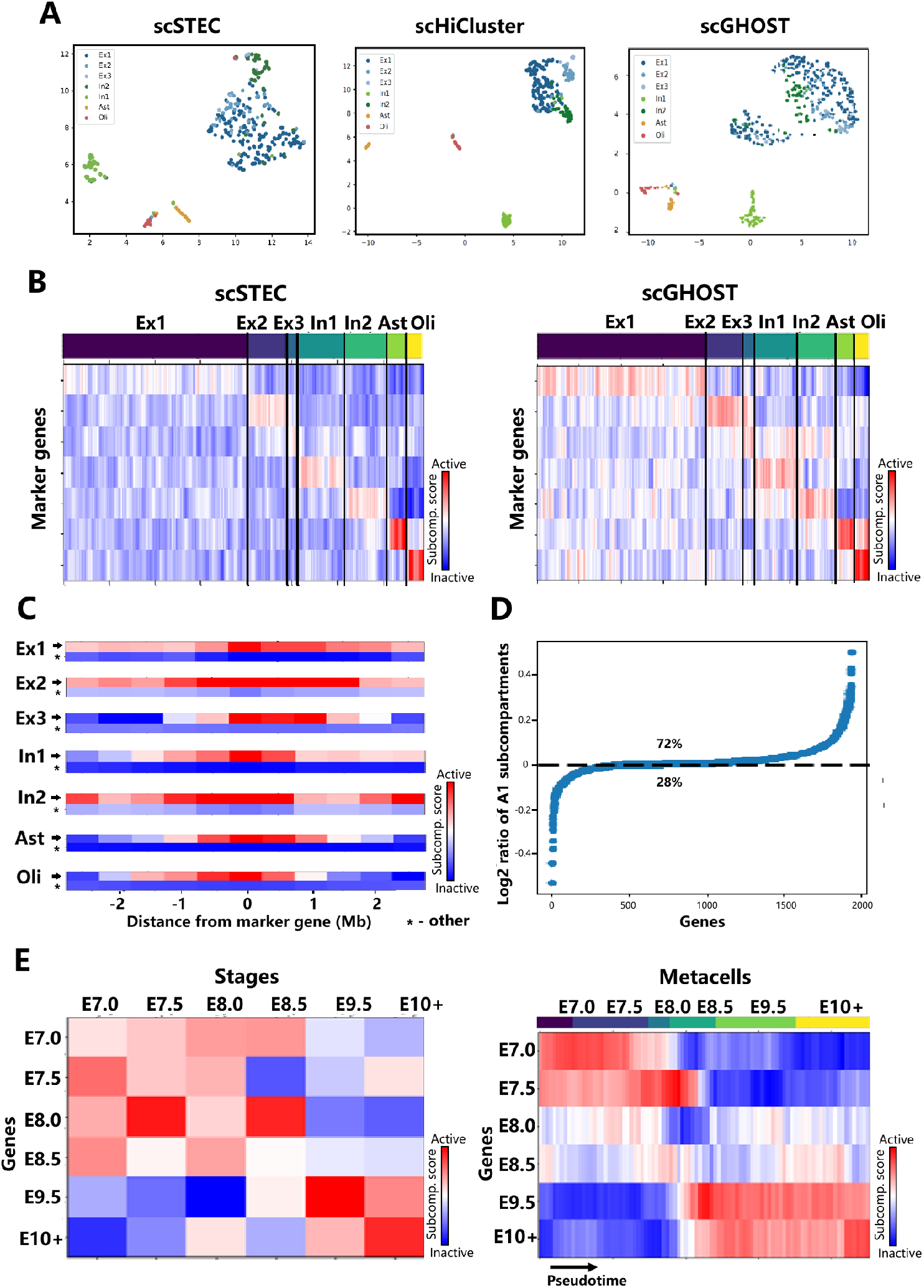
Application of scSTEC to HiRES datasets of the adult mouse cortex and developing mouse embryos. (A) UMAP embeddings derived from scSTEC subcompartment annotations, scHiCluster, and scGHOST for the adult mouse cortex. (B) Average subcompartment scores across the top 200 marker genes of each cell type in individual cells, calculated using scSTEC and scGHOST. Each column represents a single cell, and each row represents the marker-gene set of one of the seven cell types. (C) Average subcompartment scores across the 2.5 Mb flanking regions of cell-type-specific marker genes in their corresponding cell type compared with the same genomic regions in other cell types. (D) Log_2_ fold change in the percentage of A1 assignments at the TSS between actively transcribing (UMI ≥ 10) and silent (UMI = 0) cells for each gene. (E) Subcompartment dynamics during mouse embryonic development. scSTEC subcompartment scores for stage-specific marker genes across six developmental stages are shown for stage-specific pseudo-bulk Hi-C matrices (left) and metacells ordered along developmental pseudotime (right). For visualization, the metacell trajectories were smoothed over every five neighboring metacells.

To link subcompartments to gene expression, we assigned scores from 0 (B3) to 4 (A1) according to subcompartment activity and computed the mean scores for the top 200 marker genes of each cell type (Figure 4B). Across all seven cell types, marker genes tended to reside in more active subcompartments in their corresponding cell types than in other cell types. Similar cell-type-specific patterns were observed using scGHOST subcompartment annotations and scA/B values (Figure 4B and Supplementary Figure S16).

We further examined the 2.5 Mb flanking regions surrounding these marker genes. For each marker gene set, we computed the average subcompartment scores across the flanking regions in the corresponding cell type and compared them with the same genomic regions in all other cell types. The flanking regions of cell-type-specific marker genes showed significantly higher subcompartment activity in their corresponding cell types than in other cell types (Figure 4C; *P <* 6 *×* 10^−10^, Paired t-test). Moreover, although marker genes themselves consistently occupied active subcompartments, their surrounding regions showed distinct patterns of subcompartment activity across cell types. These results suggest that cell-type-specific gene expression is associated not only with local subcompartment activity at marker genes but also with cell-type-dependent differences in their surrounding chromatin environments.

We next examined the relationship between subcompartment boundaries and TAD-like domain boundaries using single-cell insulation scores (Supplementary Figure S17). We calculated single-cell insulation scores from scHiCluster-imputed contact maps at 500 kb resolution using a 2 Mb window, and aggregated the scores around subcompartment boundaries, subcompartment centers, and random genomic regions. On both chromosome 1 and the whole genome (Supplementary Figure S17), subcompartment boundaries coincided with local minima in the aggregated insulation scores, consistent with their preferential localization at TAD-like domain boundaries. For large subcompartments (*>* 2.5 Mb), insulation scores at subcompartment centers were higher than those at random genomic regions, suggesting that loci within large subcompartments were less likely to correspond to TAD-like domain boundaries.

Finally, we examined the relationship between single-cell subcompartment states and transcription within the same cell type, focusing on Ex1 cells (Figure 4D). For 72% of genes, cells actively transcribing the gene (UMI ≥ 10) were more likely to have the TSS annotated as the A1 subcompartment than cells in which the same gene was silent (UMI = 0). This pattern remained consistent when A1 and A2 were jointly considered as the A compartment (Supplementary Figure S18). These results reveal a within-cell-type association between transcriptional activity and single-cell subcompartment state.

### 3.4 scSTEC applied to developing mouse embryos

We next applied scSTEC to scHi-C data from developing mouse embryos (E7.0–E11.5) in HiRES dataset [22] to investigate subcompartment dynamics during development. We first generated single-cell RNA and scHiCluster embeddings using the HiRES data (Supplementary Figure S19). We then identified stage-specific marker genes by comparing each developmental stage with all other stages and selected the top 200 marker genes for each stage based on fold change.

At the pseudo-bulk level, we applied scSTEC to stage-specific pseudo-bulk Hi-C matrices (Figure 4E, left). Early-stage (E7.0/E7.5) marker genes preferentially occupied more active subcompartments at early developmental stages (before E8.0), whereas late-stage (E9.5/E10+) marker genes preferentially occupied more active subcompartments at later stages (E9.5/E10+). In contrast, marker genes from the intermediate stages (E8.0/E8.5) showed greater variability in subcompartment activity across development.

To examine developmental subcompartment dynamics at cellular resolution while mitigating the effects of data sparsity and cell-cycle variation, we grouped transcriptionally similar cells into metacells and averaged their independently inferred single-cell subcompartment scores at each locus, following the strategy used in our previous work [18] (Methods; Supplementary Material). We tracked subcompartment changes across developmental pseudotime. The resulting subcompartment dynamics closely recapitulated the stage-wise pseudo-bulk patterns and followed the developmental timeline (Figure 4E, right). Loci containing early-stage marker genes (E7.0/E7.5) showed high subcompartment activity early in development (before E8.0), followed by a progressive decrease. Conversely, loci containing late-stage marker genes (E9.5/E10+) transitioned toward more active subcompartments primarily after E8.5, whereas intermediate-stage marker genes (E8.0/E8.5) showed greater variability in subcompartment activity across developmental pseudotime.

We next focused on E8.5 stage-specific marker genes to examine the temporal relationship between subcompartment reorganization and gene expression in greater detail. For each gene, we constructed subcompartment and expression trajectories along developmental stages, min–max normalized each trajectory to [0, 1], and performed Louvain clustering (Supplementary Figure S20). This analysis identified five gene clusters with distinct dynamic patterns. For approximately 40% of genes (clusters 2 and 4), subcompartment changes preceded transcriptional upregulation; for 23% (cluster 1), subcompartment and transcriptional changes occurred approximately synchronously; and for 16% (cluster 5), subcompartment changes followed transcriptional activation. The remaining 21% of genes (cluster 3) showed no clear temporal correspondence between subcompartment and expression dynamics. These results revealed heterogeneous temporal relationships between subcompartment reorganization and transcriptional changes during embryonic development.

### 3.5 scSTEC applied to developing olfactory neurons

Finally, we evaluated scSTEC using LiMCA datasets, which jointly profiled 3D genome organization and transcriptomes in single cells [20]. We first analyzed the four human cell lines (GM12878, K562, eHAP, and BJ) as an additional validation of cell-type-specific subcompartment organization. Cellular embeddings based on scSTEC subcompartment annotations clearly separated the four cell types (Figure 5A). In addition, scSTEC identified a distinct cluster of metaphase cells that closely agreed with the original LiMCA annotation, with 25 of 26 cells overlapping [20]. Consistently, marker genes for each cell line tended to occupy more active subcompartments in their corresponding cell type (Figure 5A).

**Figure 5:**
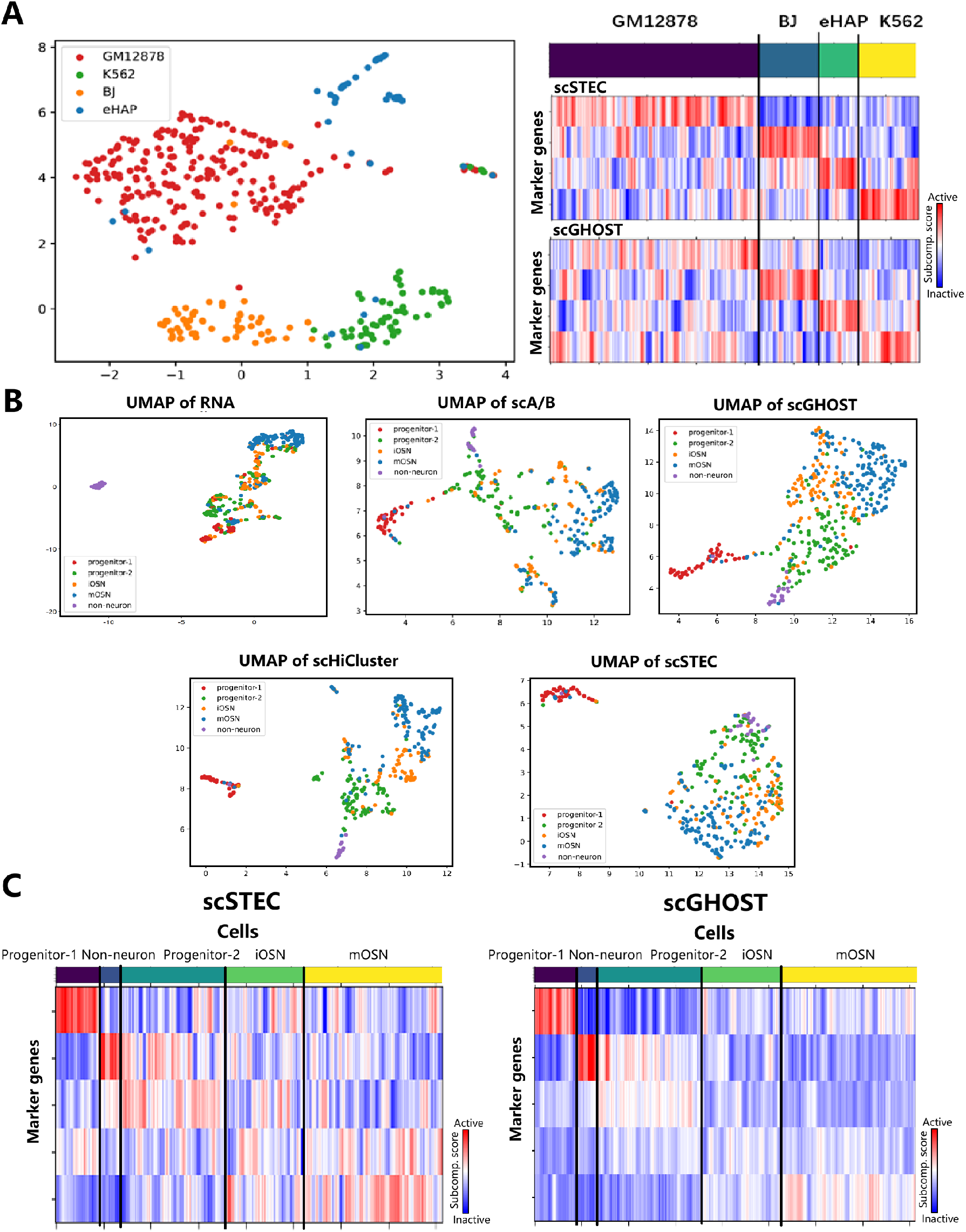
Application of scSTEC to LiMCA datasets of four human cell lines and developing olfactory sensory neurons. (A) UMAP embedding based on scSTEC subcompartment annotations for four human cell lines (GM12878, K562, eHAP, and BJ) (left), and average subcompartment scores across the top marker genes of each cell type in individual cells (right). (B) UMAP embeddings of developing olfactory sensory neurons (OSNs) derived from scRNA-seq, scSTEC subcompartment annotations, scA/B values, scHiCluster, and scGHOST. (C) Average subcompartment scores across cell-type-specific marker genes during OSN development, calculated using scSTEC and scGHOST. Cells were ordered along the developmental trajectory, and the single-cell subcompartment scores were smoothed over every five neighboring cells for visualization.

We next applied scSTEC to the LiMCA dataset of developing mouse olfactory sensory neurons (OSNs) [20], comprising 411 cells collected at six postnatal time points (days 3, 7, 14, 28, 60, and 120).

In Figure 5B, we generated cellular embeddings using scSTEC and compared them with embeddings derived from scRNA-seq, scA/B values, scHiCluster, and scGHOST. The RNA embedding revealed four major cell populations: non-neuronal cells, progenitors, immature OSNs (iOSNs), and mature OSNs (mOSNs), with progenitors, iOSNs, and mOSNs forming a developmental trajectory. The four Hi-C-based embeddings broadly recapitulated this organization but further separated the progenitors into two groups, denoted progenitor-1 and progenitor-2. Progenitor-1 was positioned outside the main Hi-C developmental trajectory, whereas progenitor-2 followed a progression toward iOSNs and mOSNs. These results revealed differences between 3D genome and transcriptional organization during OSN development, while demonstrating that scSTEC subcompartment annotations alone could resolve major cell populations and capture the principal developmental trajectory.

We next examined subcompartment assignments for marker genes of each cell type (Figure 5C). Compared with scGHOST, scSTEC showed similar enrichment of active subcompartments for marker genes in their corresponding cell types. This enrichment was less pronounced for the three cell types along the developmental lineage (progenitor-2, iOSNs, and mOSNs), consistent with their less distinct separation in the UMAP embeddings and suggesting greater sub-compartmental variability during OSN development. As an independent example, we further examined a previously identified set of mOSN-enriched genes, including well-known markers such as S100a5, Adcy3, and Gnal [27]. These genes likewise preferentially occupied more active subcompartments in mOSN cells (Supplementary Figure S21).

## 4 Discussion

In this study, we introduced scSTEC for direct chromatin subcompartment annotation from individual scHi-C matrices. Its principal contribution is to enable independent annotation of each cell without requiring dataset-wide cellular embedding, deep-learning training, or cross-cell imputation. After a common set of valid genomic bins is defined, subcompartment annotation for each cell uses only that cell’s own Hi-C matrix, without borrowing contact information from other cells. The mathematical foundation of scSTEC is the observation that the estimated inter-chromosomal matrix constructed in scGHOST has rank two, allowing the Euclidean distances between its rows to be preserved exactly in a two-dimensional representation. Building on this geometric structure, scSTEC bypasses complex upstream procedures such as deep-learning-based imputation and cellular embedding, providing a more transparent and direct analytical framework. It also substantially reduces computational cost: scSTEC was markedly faster than CPU-based scGHOST and required no GPU acceleration (Supplementary Figure S22), supporting its practical scalability to increasingly large scHi-C datasets.

Through extensive evaluations, we showed that scSTEC produced subcompartment annotations comparable to those obtained by scGHOST. On bulk Hi-C and simulated scHi-C matrices, scSTEC recovered the canonical subcompartment structures (A1, A2, B1, B2, and B3) and remained robust under severe data sparsity, with performance comparable to scGHOST even at a 1/6000 sampling rate. Across diverse real scHi-C datasets, scSTEC-derived subcompartments were consistently associated with characteristic epigenetic signatures, long-range contact patterns, cell-type-specific gene expression, and developmental dynamics. Notably, cellular embeddings constructed solely from scSTEC subcompartment annotations were sufficient to resolve major cell populations and capture developmental trajectories.

Beyond annotation, scSTEC enabled the exploration of functional associations of subcompartment organization at the single-cell level. In particular, the cell-to-cell variability of subcompartment states was associated with transcriptional variability and H3K27me3 levels, with stable loci preferentially occupying A1 and B3 subcompartments. We also found that subcompartment boundaries were preferentially associated with TAD-like domain boundaries, suggesting a relationship between these two levels of chromatin organization. Moreover, cell-type-specific marker genes tended to occupy more active subcompartments in their corresponding cell types. Together, these findings illustrate how individual-cell subcompartment annotations can be used to investigate the relationships between 3D genome organization, epigenetic state, and transcriptional heterogeneity.

Several limitations and future directions warrant consideration. First, scSTEC requires scHi-C data with sufficient informative contacts; in low-coverage datasets, a substantial fraction of cells may fail the quality-control criteria and therefore cannot be reliably analyzed. Second, scSTEC uses CpG density to identify the initial active and inactive reference loci. Although this strategy performed consistently across the datasets examined here, its generalizability to broader cellular contexts remains to be systematically evaluated. Incorporating additional genomic features may provide more flexible reference selection in future extensions. Finally, although we used matched transcriptomic and epigenetic measurements to characterize the biological associations of scSTEC-derived subcompartments, these modalities are not incorporated into the inference itself. Future extensions could jointly model scHi-C with other single-cell omics modalities to investigate the relationships between subcompartment organization, epigenetic state, and transcriptional regulation.

## Supporting information

Supplementary methods, results and figures

## Acknowledgments

We thank Zhiyuan Liu and Dong Xing for helpful discussions concerning real data analysis. We thank Jian Ma and members of his group for helpful clarifications regarding technical details of scGHOST.

## Author contributions

H.G. conceived and supervised the project. Y.P. and L.Y. designed and implemented the scSTEC algorithm and performed the simulation, benchmarking, and real-data analyses. Y.P., L.Y., and H.G. wrote the manuscript. All authors read and approved the final manuscript.

## Supplementary data

Supplementary data are available at Bioinformatics online.

## Conflict of interests

None declared.

## Funding

H.G. was supported by National Natural Science Foundation of China (T2225001) and National Key *R&D* Program of China (2023YFF1204700).

## Data availability

The source code is available at https://github.com/Ge-lab-pku/scSTEC. The source code is also archived on Zenodo and can be found at https://doi.org/10.5281/zenodo.23750594. All data used are publicly available online; details can be referred to in the supplementary data.

## Notes

### Competing Interest Statement

The authors have declared no competing interest.

