## Supplementary methods, results and figures for "scSTEC: Annotating Chromatin Subcompartments from Individual Single-Cell Hi-C Data via Direct Two-Dimensional Embedding and Chromosome-wise Consensus"

### 1 Supplementary Methods

#### 1.1 Bulk Hi-C subcompartment annotation

For bulk Hi-C data, we used the O/E-normalized (observed/expected) Hi-C contact matrix and A/B compartment information derived from bulk PCA to compute the 2D genomic representation $[\vec{a}, \vec{b}]$. Hi-C matrices were handled using `cooler` [1], whereas O/E normalization and bulk PCA were performed using `cooltools` [2].

For scHi-C data, in contrast, we directly used raw contact maps because O/E normalization is less reliable under extreme data sparsity. Following the idea of scA/B values, CpG density was used as a proxy for A/B compartment identity when constructing the 2D genomic representation. The subsequent 2D embedding, clustering, labeling, and chromosome-wise consensus procedures were performed as described in the main Methods.

#### 1.2 Data cohorts and accessions

1. **GM12878 datasets**: Bulk Hi-C maps from [3] were obtained from GEO under accession GSE63525. Deep single-cell Micro-C matrices [4] were retrieved from GEO Superseries GSE281150. We used the scMicro-C subseries GSE192759, comprising 355 cells. Single-cell Hi-C contact maps from [5] were obtained from the published repository (`https://noble.gs.washington.edu/proj/schic-topic-model`) at 500 kb resolution.

   Epigenetic data for GM12878 were obtained from ENCODE: ENCFF167NBF (H3K27me3), ENCFF171MDW (H3K36me3), ENCFF803DJF (H3K79me2), ENCFF776OVW (H3K9me3), ENCFF818GNV (H3K4me3) and ENCFF180LKW (H3K27ac). Repli-seq data were also obtained from ENCODE: ENCFF001GNK (G1), ENCFF001GNR (S1), ENCFF001GNT (S2), ENCFF001GNX (S3), ENCFF001GOA (S4), and ENCFF001GNN (G2).

2. **LiMCA datasets**: The LiMCA dataset from [6] was obtained from GEO (GSE240128). Both the four-cell-line dataset (GM12878, K562, eHAP, and BJ, 389 cells) and the developing olfactory sensory neuron (OSN) dataset (411 cells) were used for analysis. The 100 marker genes for mOSNs were obtained from [7].

3. **HiRES datasets**: The HiRES dataset from [8] was obtained from GEO (GSE223917). Both the adult mouse cortex dataset (399 quality-filtered cells) and the developing mouse embryo dataset (7,469 total cells across stages E7.0–E11.5) were used for analysis. Developmental analyses focused on the 3,247 cells constituting the main neural and mesenchymal lineages. Cell-type annotations were provided with the original dataset and were based on the matched scRNA-seq data.

4. **CHARM dataset**: The CHARM dataset from [9] was obtained from GEO (GSE303006). The dataset profiling H3K27ac in the adult mouse cortex was used for analysis. To eliminate cell-type confounding, analyses were restricted to 538 L3/4 IT excitatory neurons. Cell-type annotations were provided with the original dataset.

#### 1.3 Preprocessing of real scHi-C and scRNA-seq data

For all mouse datasets, genomic coordinates and gene annotations were based on the GRCm38 reference genome with GENCODE M23 annotations. For human datasets, GRCh37 was used for GM12878 from [5], whereas GRCh38 was used for LiMCA and scMicro-C. Human gene annotations were obtained from the UCSC Genome Browser [10]. CpG density profiles were computed by quantifying CG dinucleotide frequencies in the corresponding reference genomes.

We used `cooler` [1] and `cooltools` [2] to perform O/E normalization, matrix balancing, and compartment annotation for bulk and pseudo-bulk Hi-C matrices. Our analyses were restricted to autosomes (chr1–19 for mouse and chr1–22 for human). Unless otherwise specified, Hi-C matrices were binned at 500 kb resolution.

###### Rao's bulk data

We first performed KR normalization on both intra- and inter-chromosomal Hi-C matrices using the KR normalization factors provided with the Rao dataset [3]. For the intra-chromosomal matrices, we then used `cooler` and `cooltools`, as described above, to perform O/E normalization and bulk PCA.

Rao's subcompartment annotations were reproduced using a Gaussian hidden Markov model (GHMM) applied to inter-chromosomal contact matrices. Following Rao et al. [3], odd-numbered chromosomes were arranged along one dimension and even-numbered chromosomes along the other, and the GHMM analysis was performed in both directions. Before fitting the GHMM, the inter-chromosomal contact data were log-transformed and z-score normalized, following the procedure of Rao et al. [3]. The GHMM was implemented using the `hmmlearn` package.

##### LiMCA

For the analysis in Figure 3, we used only GM12878 cells from the LiMCA dataset. For Figure 5, we used all five cell populations, with five cells from the four human cell lines excluded during quality control. Metaphase cells were not excluded but formed a distinct cluster in the cellular embedding. Cell-type annotations and cell-cycle phase labels were obtained from the original dataset.

##### scRNA-seq

Because gene-expression variance in single-cell RNA-seq data is strongly dependent on mean expression [11], the raw standard deviation tends to assign greater variability to highly expressed genes. We therefore used the coefficient of variation (CV) as a normalized measure of transcriptional variability. For each gene, the CV was calculated across all single cells as the ratio of the standard deviation to the mean, $\mathrm{CV} = \sigma/\mu$.

#### 1.4 Variability of subcompartments

Because subcompartment annotations are discrete labels, we quantified their cell-to-cell variability using the entropy of the subcompartment distribution at each genomic locus.

Specifically, for genomic locus $j$ we defined the subcompartment variability as

$$V_j = -\sum_{m=0}^{4} P_{mj} \log P_{mj},$$

where $P_{mj} = \frac{1}{N} \sum_{i=1}^{N} \mathbf{1}_{L_i(j)=m}$ is the proportion of cells assigned to subcompartment $m$ at locus $j$ and $L_i(j)$ is the subcompartment label at locus $j$ in cell $i$ as defined in Section 2.7 in the main text, with $0 \log 0$ defined as 0.

A low value of $V_j$ indicates that most cells share the same subcompartment label at locus $j$ and therefore corresponds to a stable locus. In contrast, a high value indicates that the subcompartment labels are more broadly distributed across cells and therefore corresponds to a variable locus.

#### 1.5 Comparison of intra- and inter-subcompartment contacts

For the analyses in Figures 2C and 3A, we computed bulk and pseudo-bulk observed-over-expected (O/E) contact maps. For each subcompartment, contact pairs were divided into intra- and inter-subcompartment groups. The intra-subcompartment group comprised contacts for which both genomic regions were assigned to the focal subcompartment, whereas the inter-subcompartment group comprised contacts between the focal subcompartment and all other subcompartments.

We compared the O/E contact values between the intra- and inter-subcompartment groups using a two-sample t-test implemented with the `ttest_ind` function in SciPy [12].

#### 1.6 Cellular embedding

All cellular embeddings were generated using UMAP.

1. **Transcriptomic embeddings**. Single-cell RNA-seq profiles were normalized using `SCTransform` with default parameters in Seurat, and 3,000 highly variable genes were selected. Principal component analysis (PCA) was then performed, and the leading 25 principal components were used for UMAP embedding. Graph-based clustering was performed using Seurat's `FindClusters` with `resolution` = 1.5.

2. **Subcompartment embeddings (scSTEC and scGHOST)**. Single-cell subcompartment profiles, encoded as discrete values from 0 to 4 corresponding to B3, B2, B1, A2, and A1, respectively, were used for cellular embedding. For each chromosome, the subcompartment profiles were first reduced to 20 PCs. The resulting PCs were concatenated across chromosomes and subjected to a second PCA, from which 20 PCs were retained. These 20 PCs were used for UMAP embedding with the Python `umap-learn` package (`n_neighbors`=10, `min_dist`=0.0; all other parameters were set to default values).

3. **scA/B compartment embeddings**. Rank-normalized scA/B values, scaled to $[0, 1]$, across all autosomes were reduced to 15 PCs and then used for UMAP embedding with the same UMAP parameters as above.

4. **scHiCluster imputation embeddings**. Imputed single-cell contact matrices generated by scHiCluster were flattened, retaining only contacts between genomic loci separated by less than 10 Mb. Following the original protocol [13], 50 PCs were computed for each chromosome. These PCs were concatenated across chromosomes and further reduced to 20 PCs, which were then used for UMAP embedding with the same parameters as above.

#### 1.7 Differential expression analysis and marker gene selection

Cell-type- and stage-specific transcriptional markers were identified from normalized single-cell RNA-seq data using Seurat [14], following a procedure similar to that used in our previous work [15]. Unless otherwise specified, positive marker genes were identified using `FindAllMarkers(only.pos = TRUE)` with default parameters.

1. **Adult mouse cortex (HiRES)**: Using the cell-type annotations provided with the original dataset, we identified up-regulated genes for each of the seven brain cell types—three excitatory neuron clusters (Ex1–3), two inhibitory neuron clusters (In1–2), astrocytes (Ast), and oligodendrocytes (Oli)—relative to all remaining cells. For each cell type, the top 200 marker genes ranked by average $\log_2$ fold change were retained for downstream subcompartment analyses.

2. **Developing mouse embryos (HiRES)**: To identify stage-specific transcriptional programs, cells were grouped by embryonic sampling time point (E7.0, E7.5, E8.0, E8.5, E9.5, and E10+), irrespective of lineage. For each developmental stage, enriched genes were identified relative to cells from all remaining stages, and the top 200 positive markers ranked by average $\log_2$ fold change were retained.

3. **LiMCA datasets**: Marker genes for the four human cell lines and the five olfactory sensory neuron populations (progenitor-1, progenitor-2, immature OSNs, mature OSNs, and non-neuronal cells) were identified using the same differential expression procedure, with the top 200 enriched marker genes retained for each group. For mature OSNs, we additionally included the 100 established olfactory receptor-associated marker genes reported in [7].

#### 1.8 Embryonic Metacell Construction and Pseudotime inference

To smooth downstream subcompartment trajectories and reduce cell-to-cell fluctuations while preserving regulatory heterogeneity during mouse embryogenesis, we constructed metacells by grouping transcriptionally similar single cells from the HiRES embryo dataset, following the procedures in [8, 15]. Mitotic (M-phase) cells were excluded, leaving 3,217 cells. Single-cell RNA-seq profiles were normalized using `SCTransform`, and cells were clustered using Louvain community detection based on the top 25 principal components, with a clustering resolution of 35. Clusters containing fewer than five cells were excluded, resulting in 167 metacells with a median size of 18 cells.

For each metacell, the discrete subcompartment labels (encoded from 0 to 4) of its constituent single cells were averaged at each genomic locus to obtain a continuous metacell-level subcompartment score. For transcriptomic analysis, metacell expression profiles were obtained by averaging the expression profiles of their constituent cells, followed by scaling using `ScaleData`, and PCA was performed using the top 2,000 variable features. The cell type of each metacell was assigned by majority vote among its constituent single cells.

The transcriptomic trajectory of mouse embryogenesis comprises two prominent differentiation lineages: an early neural ectoderm (EN) trajectory and a mixed late mesenchyme (MLM) trajectory, both diverging from early epiblast/primitive streak (EPI) progenitors [8]. To examine temporal subcompartment dynamics, we performed pseudotime inference for the EN trajectory

(2,296 cells) using `Monocle3` [16], based on the transcriptomic embedding before metacell construction, following the procedures in [12, 15]. EPI progenitors were designated as the root for pseudotime ordering. The pseudotime of each metacell was then calculated as the mean pseudotime of its constituent cells.

### 2 Supplementary Results

#### 2.1 Subcompartment annotation at higher resolutions

Increasing resolution (i.e., decreasing bin size) reduces the number of contacts per bin and increases data sparsity, similar to the effect of downsampling. We therefore performed downsampling experiments on high-coverage single-cell datasets to establish empirical criteria for annotation reliability and cross-resolution consistency (Supplementary Figure S14). Specifically, we downsampled scMicro-C cells at rates ranging from 1/2 to 1/12 and LiMCA cells at rates ranging from 1/2 to 1/40. At each rate, downsampling was independently repeated 50 times using the same cell.

For each downsampled sample, we evaluated annotation concordance with the full-depth annotation using the intersection score defined in the main text. At the most extreme downsampling rates (1/12 for scMicro-C and 1/40 for LiMCA), more than 80% of samples failed quality control. Intersection scores decreased with increasing downsampling and approached a low-level plateau beyond sampling fractions of 1/5 for scMicro-C and 1/12 for LiMCA, with no significant differences from those at the most extreme downsampling rates ($p > 0.05$). We therefore selected 1/5 and 1/12, respectively, as empirical thresholds for reliable annotation. Based on the intersection scores at these thresholds, we adopted a minimum intersection score of 0.25 as a practical criterion for cross-resolution annotation consistency.

Separately, we examined cell retention rates ($1-$proportion of filtered samples) across the downsampling experiments. The fraction of samples failing quality control increased sharply around 1/10 for scMicro-C and 1/35 for LiMCA, reaching approximately 25%. We therefore adopted a maximum QC failure fraction of 0.25 (equivalently, a cell retention rate of at least 0.75) as a separate empirical criterion for real-data applications.

Together, the annotation concordance and cell retention criteria suggested that scSTEC could reliably annotate subcompartments at resolutions of approximately 50–100 kb in recent high-coverage datasets such as LiMCA and scMicro-C (Supplementary Figure S15). These thresholds are empirical guidelines that depend on data coverage and sequencing depth and may require adjustment for other datasets.

doi: 10.1093/bioinformatics/btz540. URL `https://doi.org/10.1093/bioinformatics/btz540`.

### 3 Supplementary Figures

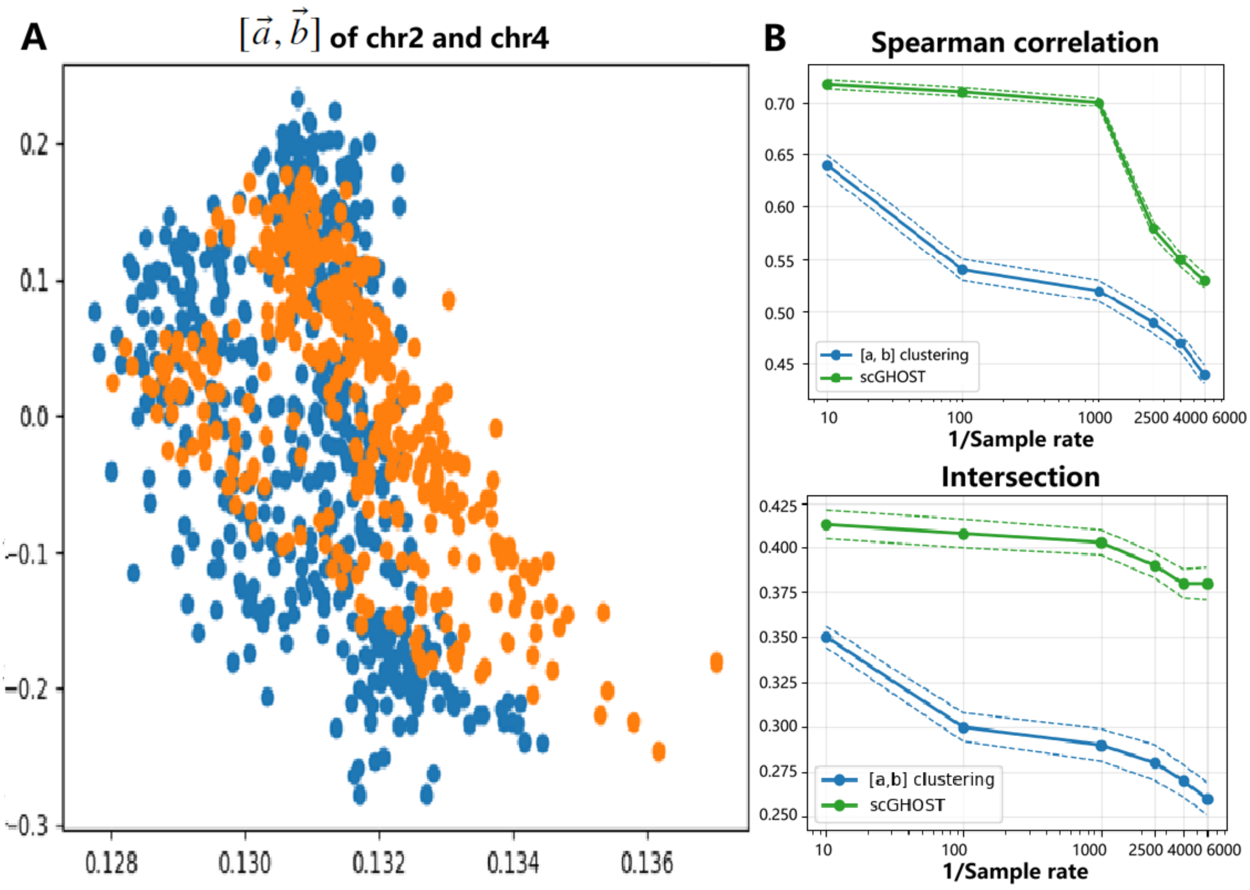


**Supplementary Fig. S1. The 2D representation $[\vec{a},\ \vec{b}]$ in scGHOST.**

(A) Visualization of $[\vec{a},\ \vec{b}]$ for chromosomes 2 and 4.

(B) Direct clustering of $[\vec{a},\ \vec{b}]$ showed lower agreement with Rao's subcompartment annotations than the scGHOST framework, as measured by Spearman correlation and intersection. Rao's bulk Hi-C data were downsampled at rates from 1/10 to 1/6000 to generate 50 simulated scHi-C datasets.

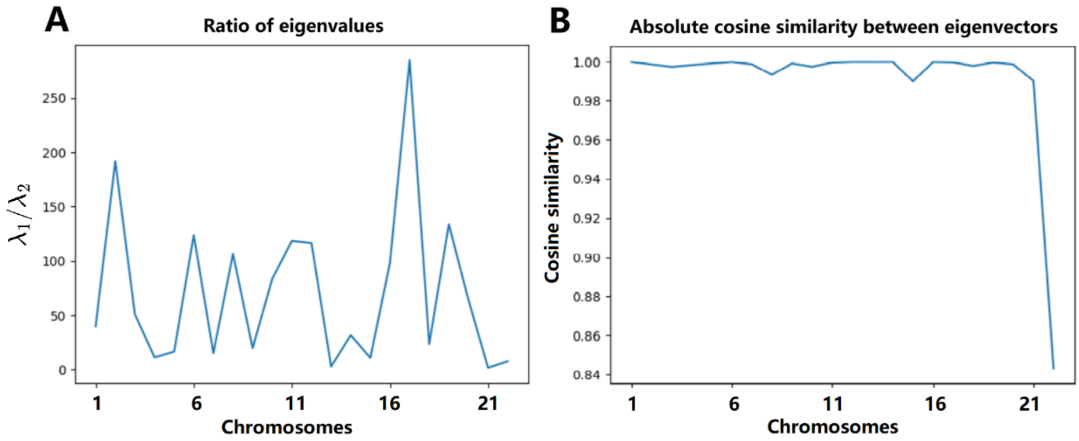


**Supplementary Fig. S2: Eigenvalues and eigenvectors used in the construction of the 2D genomic embedding from Rao's bulk Hi-C data.**

(A) Ratio of the two eigenvalues, $\lambda_1/\lambda_2$, across chromosomes, where $\lambda_1$ and $\lambda_2$ denote the larger and smaller eigenvalues, respectively. The eigenvalue ratio varied across chromosomes, reflecting different relative contributions of the two embedding directions. (B) Absolute cosine similarity between eigenvectors across chromosomes, using chromosome 1 as the reference. Both eigenvector directions were highly consistent across chromosomes.

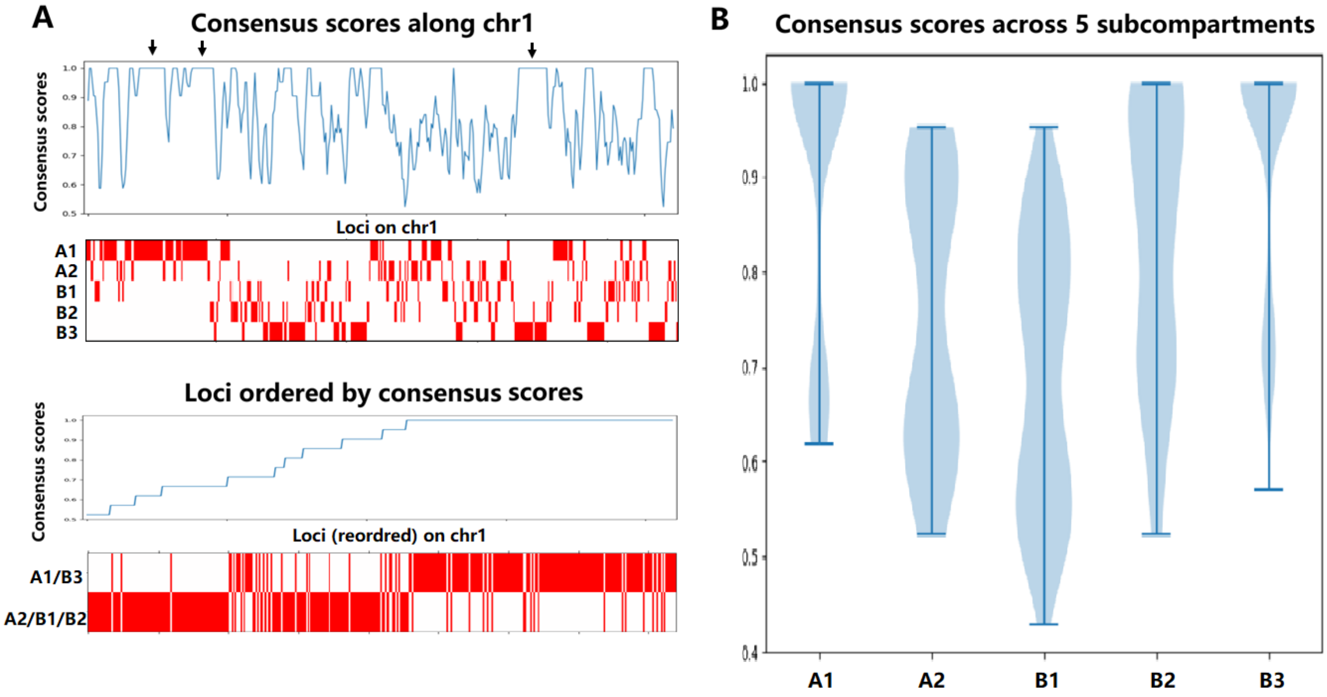


**Supplementary Figure S3. Consensus scores for Rao's bulk Hi-C data.**
(A) Consensus scores along chromosome 1. Regions with high consensus scores are indicated by black arrows and are predominantly associated with the A1 and B3 subcompartments. This pattern is further illustrated by reordering genomic loci according to their consensus scores.
(B) Comparison of consensus scores across the five subcompartments using all chromosomes.

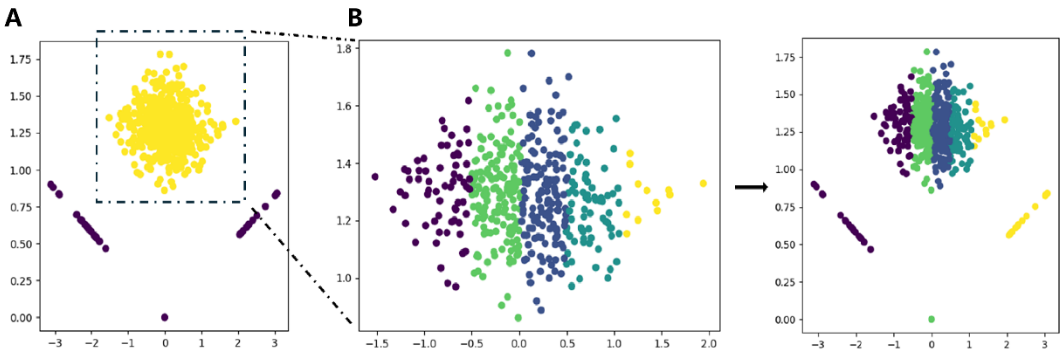


**Supplementary Fig. S4. Visualization of the 2D genomic embedding for an individual scMicro-C cell.**

(A) 2D embedding of genomic loci on chromosome 1 using chromosome 2 as the reference chromosome. Yellow points indicate loci retained for clustering, whereas purple points indicate loci excluded from the initial clustering. If fewer than 20% of the loci were retained for clustering, the corresponding cell was excluded from downstream analysis.

(B) Left: clustering of the retained loci shown in (A). Right: clustering result after assigning the initially excluded loci to their nearest clusters based on Euclidean distance in the 2D embedding space.

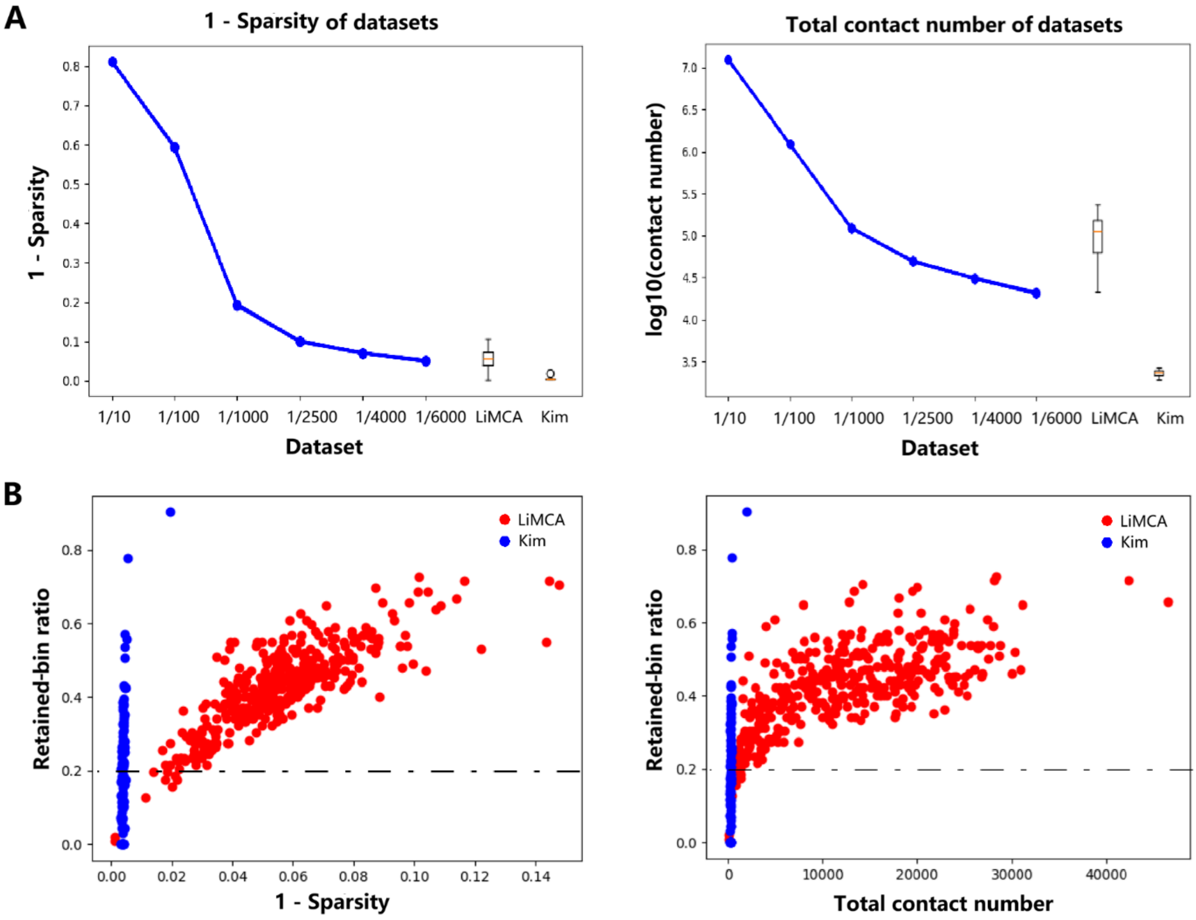


**Supplementary Figure S5. Comparison of data quality between downsampled and real scHi-C datasets.**

The LiMCA and Kim datasets were used for comparison. All datasets were derived from the human GM12878 cell line.

(A) 1-sparsity and total contact number of the downsampled and real datasets. The downsampled data were comparable to the LiMCA data in terms of these two measures and showed substantially higher data coverage than the Kim data.

(B) Relationship between the retained-bin ratio, 1-sparsity, and total contact number. The retained-bin ratio was defined as the proportion of genomic bins remaining after the scSTEC quality-control procedure. Each point represents one cell, and we only plotted the minimum values across all chromosomes for each cell. Based on the LiMCA dataset, we empirically excluded cells with 1-sparsity<0.02 (sparsity>0.98) or fewer than 3,000 contacts.

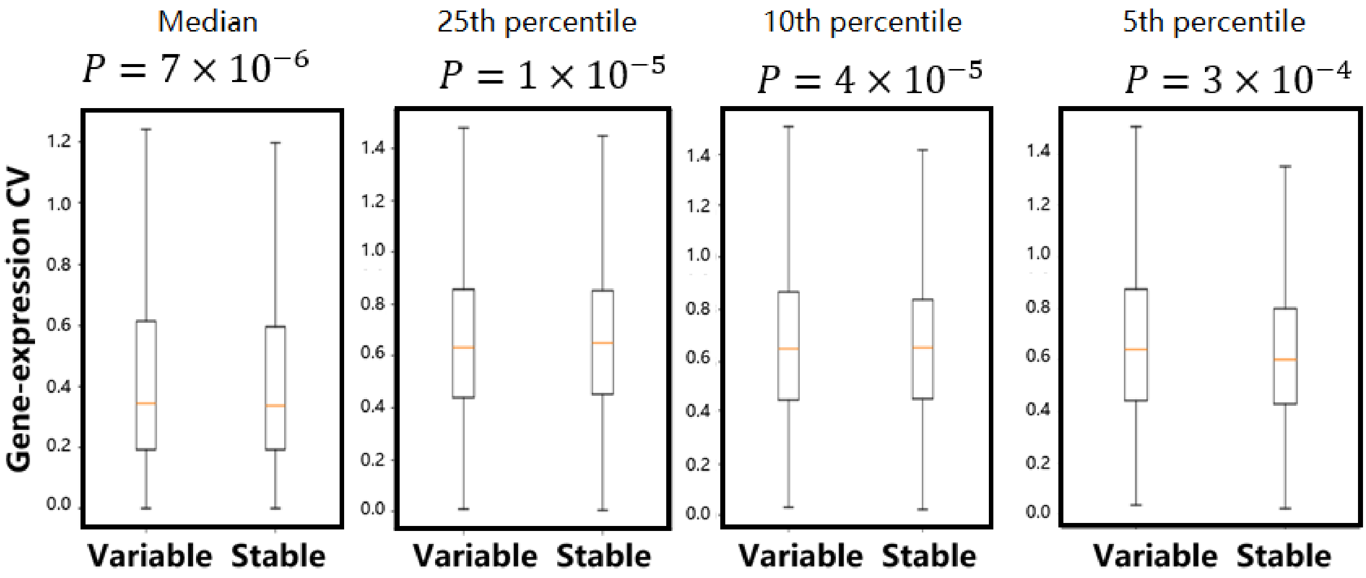


**Supplementary Fig. S6. Robustness of the identification of variable and stable loci.** We compared gene-expression CV between variable and stable loci using the top and bottom 50%, 25%, 10%, and 5% of subcompartment variability, respectively, in Ex1 cells from the HiRES dataset. All p-values were calculated using one-sided t-tests.

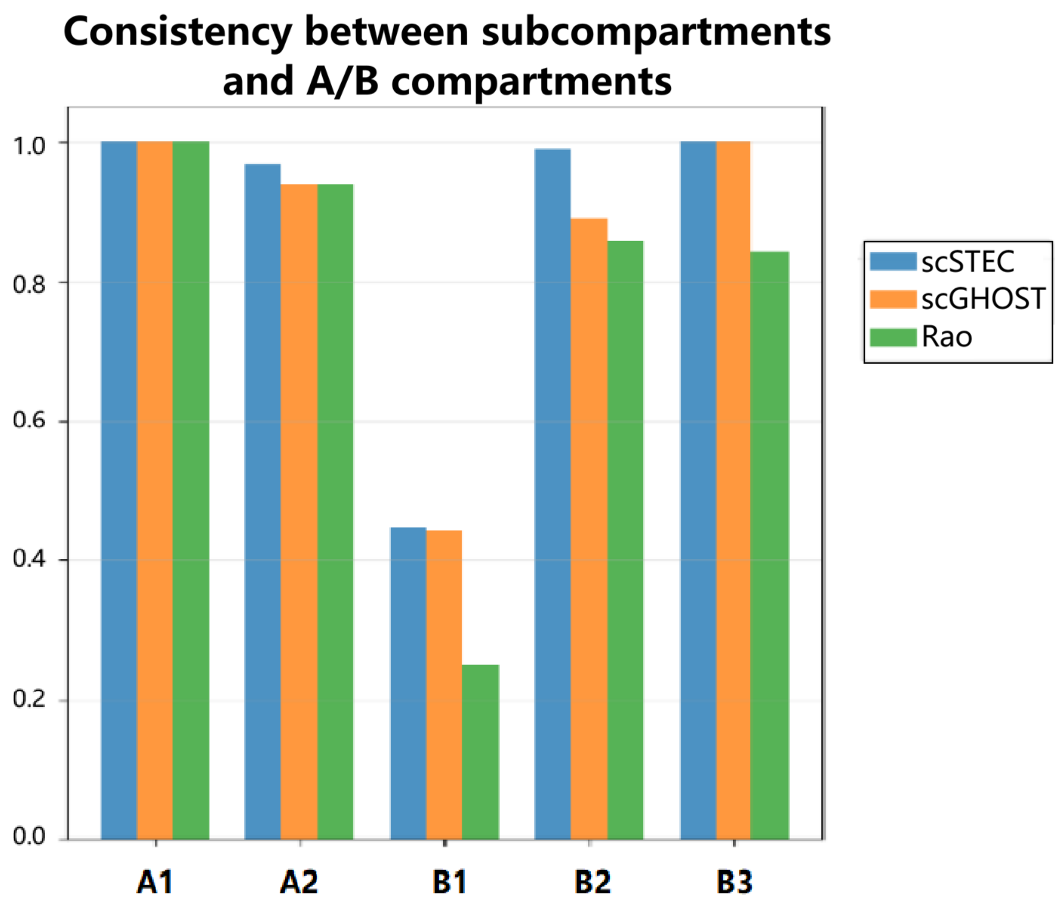


**Supplementary Fig. S7. Consistency between subcompartment annotations and A/B compartments.**

We compared subcompartment annotations with binary A/B compartment annotations derived from PCA of bulk Hi-C data. For each subcompartment, consistency was defined as the proportion of loci assigned to the corresponding A/B compartment: the proportion of A-compartment loci for A1 and A2, and the proportion of B-compartment loci for B1, B2, and B3. scSTEC and scGHOST showed high consistency with the A/B compartment annotations across the five subcompartments, whereas Rao's annotation generally showed lower consistency.

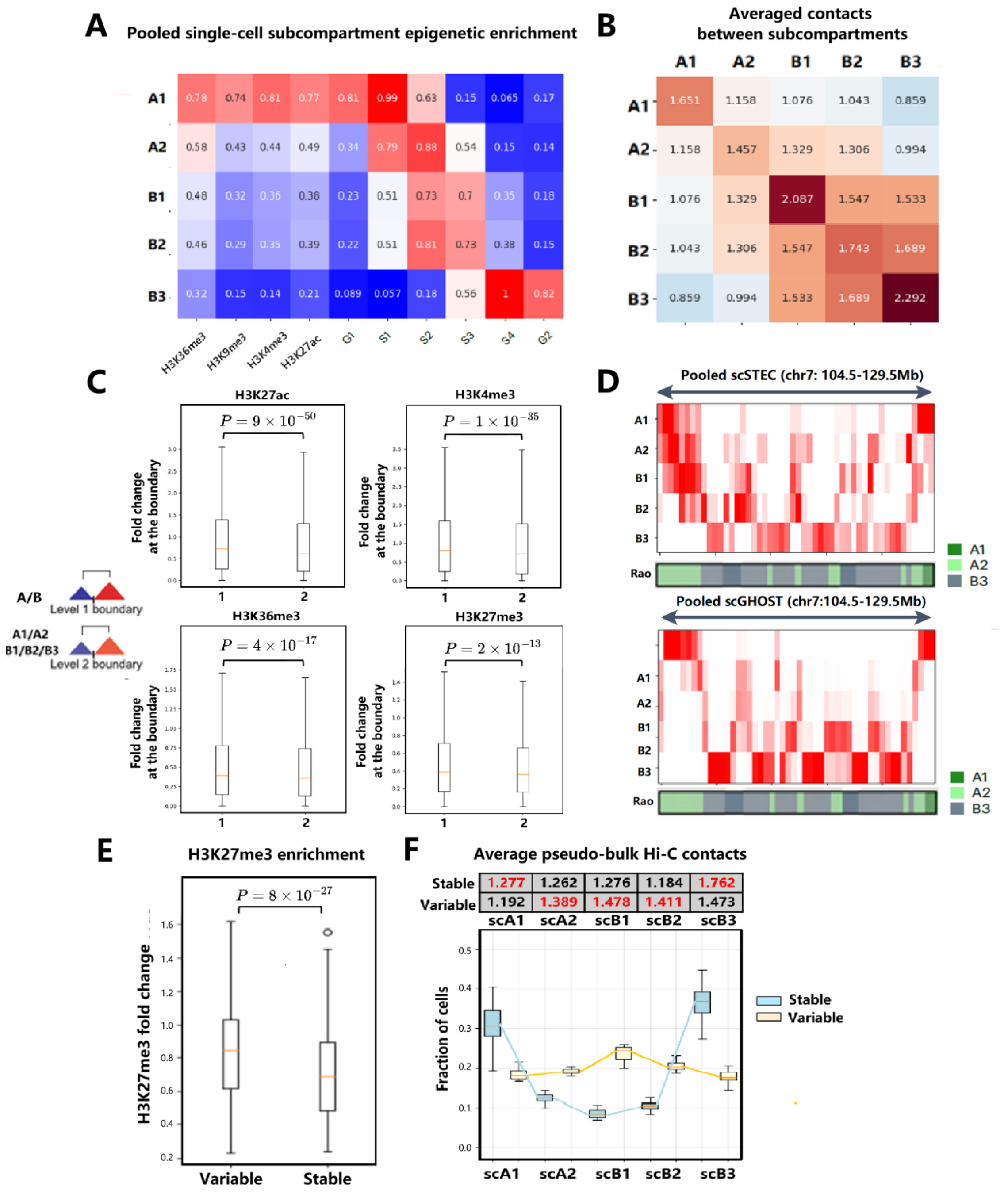


**Supplementary Figure S8. Analysis of the Kim GM12878 scHi-C dataset.**

More than 50% of cells in this dataset were excluded by the scSTEC quality-control procedure, and the remaining cells were used for the following analyses.

(A) Enrichment of epigenomic signals across pooled single-cell subcompartments.

(B) Averaged pseudo-bulk contacts among pooled single-cell subcompartments.

(C) Fold changes of H3K27ac, H3K4me3, H3K36me3, and H3K27me3 at level 1 and level 2 boundaries for each cell.

(D) Visualization of pooled single-cell subcompartment annotations obtained using scSTEC and scGHOST.

(E) H3K27me3 enrichment at genomic regions with variable and stable subcompartment annotations.

(F) Top: Average pseudo-bulk Hi-C contacts between stable/variable genomic regions and each pooled single-cell subcompartment. Bottom: Fractions of single-cell subcompartment annotations at stable (blue) and variable (yellow) genomic loci across all chromosomes.

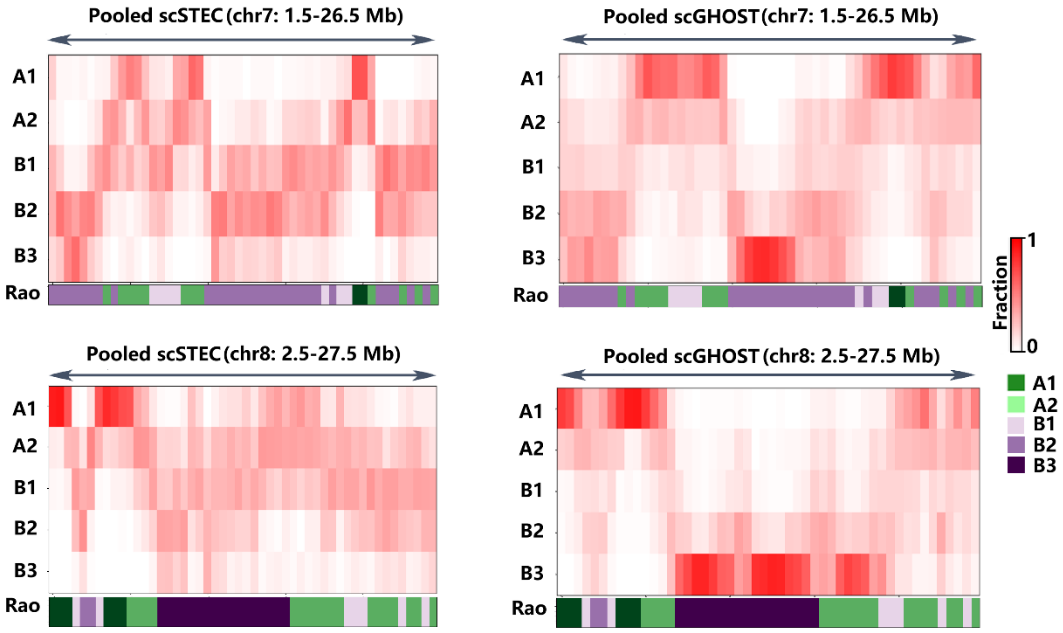


**Supplementary Fig. S9. Visualization of pooled single-cell subcompartments and bulk subcompartments in two additional 25-Mb regions on chromosomes 7 and 8.** Pooled single-cell subcompartment annotations obtained using scSTEC showed strong agreement with Rao's bulk subcompartment annotations and were visually comparable to those obtained using scGHOST.

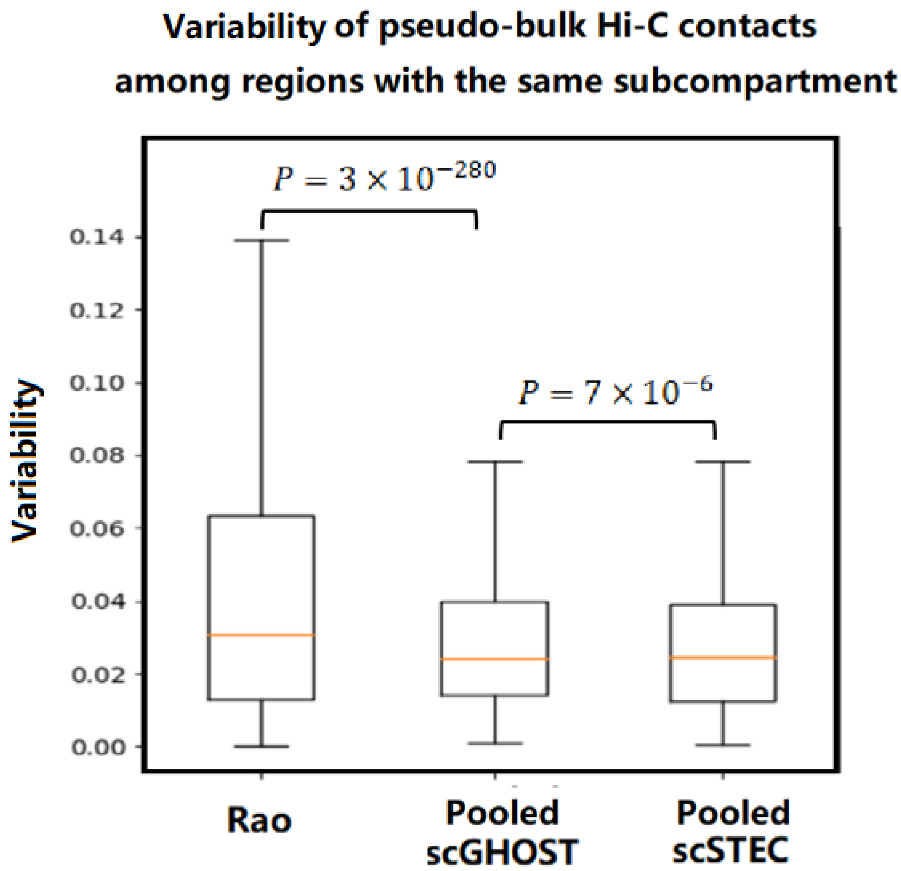


**Supplementary Figure S10. Pooled single-cell subcompartments show greater consistency with pseudo-bulk Hi-C contact patterns.**

Pseudo-bulk Hi-C contacts between each pair of subcompartments showed lower variability when loci were grouped according to the pooled single-cell subcompartment annotations from scSTEC or scGHOST than when grouped according to Rao's bulk subcompartment annotations.

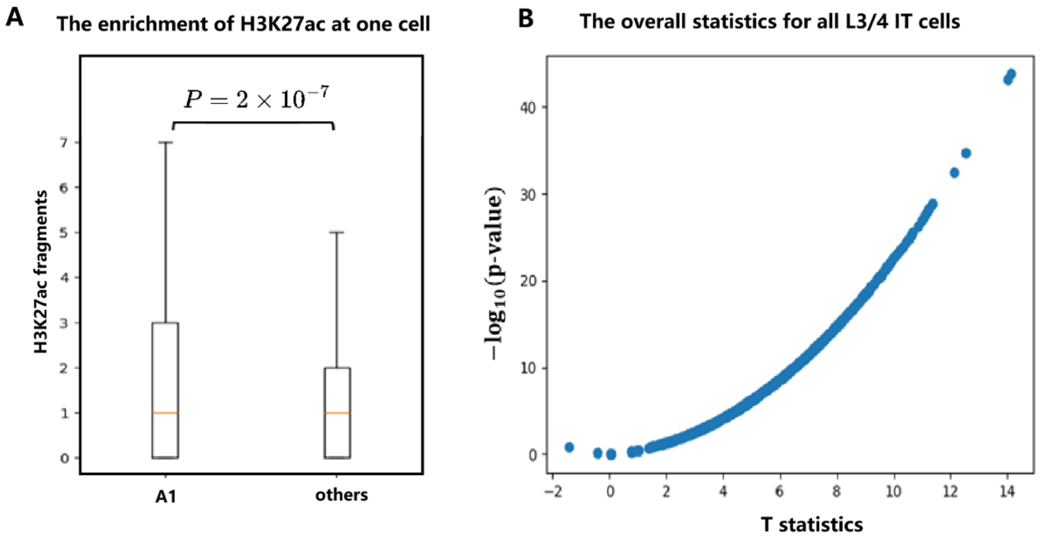


**Supplementary Figure S11. Single-cell H3K27ac enrichment across subcompartments in the CHARM dataset.**

(A) An example illustrating H3K27ac enrichment in the A1 subcompartment of an individual cell.

(B) Cell-wise comparison of H3K27ac enrichment between A1 and all other subcompartments combined using t-tests, following the procedure illustrated in (A). Each point represents one cell. The x-axis shows the t-statistic and the y-axis shows the p-value.

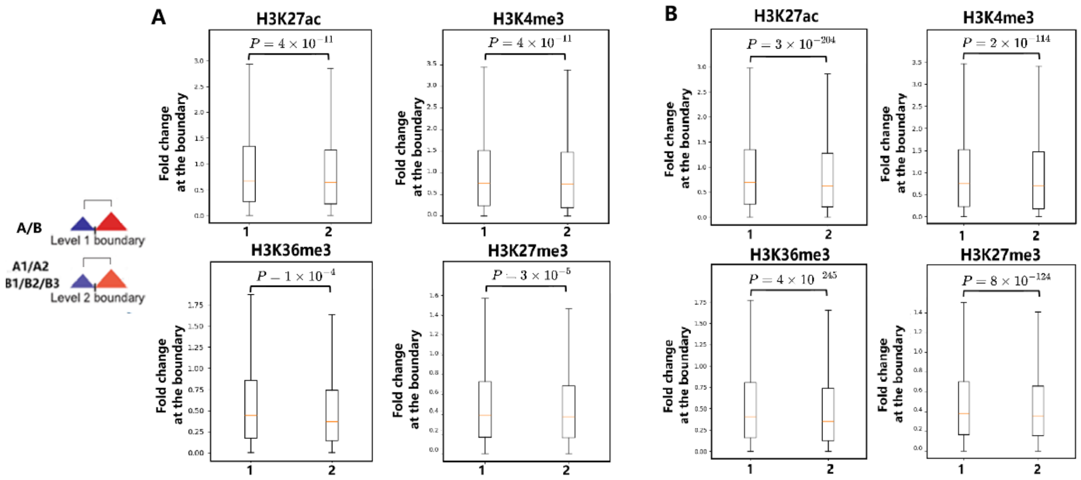


**Supplementary Figure S12. Epigenetic differences at subcompartment boundaries for pooled and single-cell subcompartments.**

Fold changes of H3K27ac, H3K4me3, H3K36me3, and H3K27me3 at level 1 and level 2 subcompartment boundaries for (A) pooled single-cell subcompartments and (B) individual-cell subcompartments in the human GM12878 cell line. For the single-cell analysis, fold changes were calculated separately for each cell.

**Enrichment of pseudo-bulk contacts**

| | scA1 | scA2 | scB1 | scB2 | scB3 |
|---|---|---|---|---|---|
| Stable | 1.864 | 1.213 | 1.135 | 1.241 | 1.753 |
| Variable | 1.690 | 1.419 | 1.470 | 1.384 | 1.441 |

**Supplementary Fig. S13. Averaged pseudo-bulk Hi-C contacts between stable/variable genomic regions and each pooled single-cell subcompartment.**

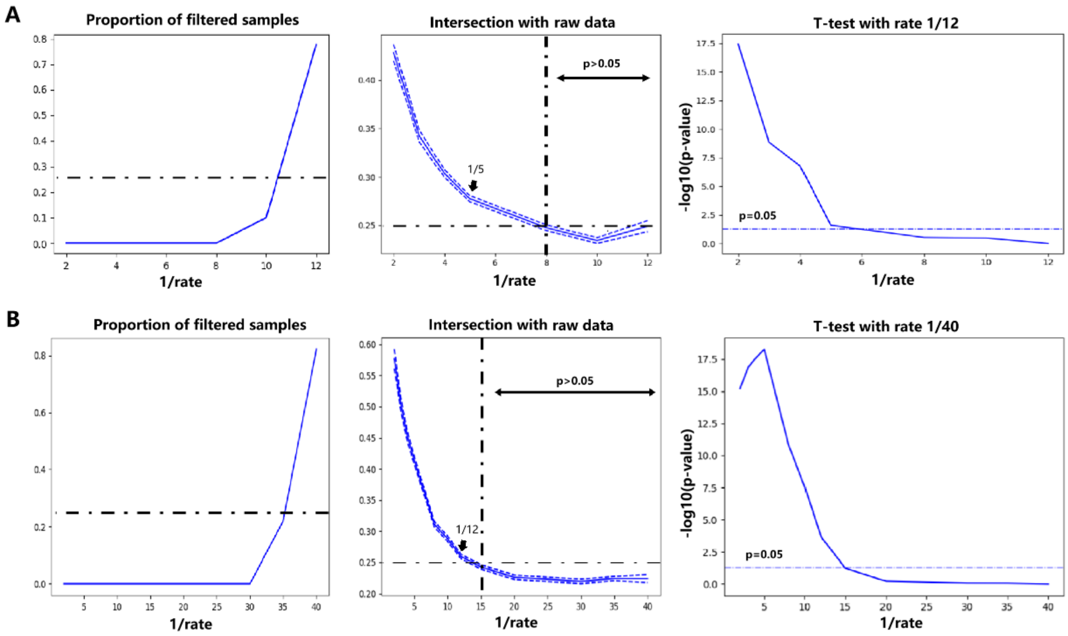


**Supplementary Figure S14. Downsampling of high-coverage scHi-C data for empirical threshold selection.**

We downsampled (A) one scMicro-C cell at rates of 1/2, 1/3, 1/4, 1/5, 1/8, 1/10 and 1/12, and (B) one LiMCA cell at rates of 1/2, 1/3, 1/4, 1/5, 1/8, 1/10, 1/12, 1/15, 1/20, 1/30, 1/35, and 1/40. At each downsampling rate, the experiment was independently repeated 50 times.

For each downsampling rate, we calculated the proportion of samples that failed the scSTEC quality-control procedure and the intersection between the subcompartment annotations obtained from the downsampled and corresponding full-depth data. In the right panels, intersection values at each downsampling rate were compared with those at the reference low-coverage rate (1/12 for scMicro-C and 1/40 for LiMCA) using t-tests, with P>0.05 used to identify rates showing no significant difference from the low-coverage reference. Based on these analyses, we empirically selected 1/5 for scMicro-C and 1/12 for LiMCA as practical lower bounds for reliable downsampling. Together with the observed increase in the proportion of samples failing quality control, these results motivated empirical thresholds of 0.25 for intersection and 0.25 for the proportion of filtered samples in subsequent real-data analyses.

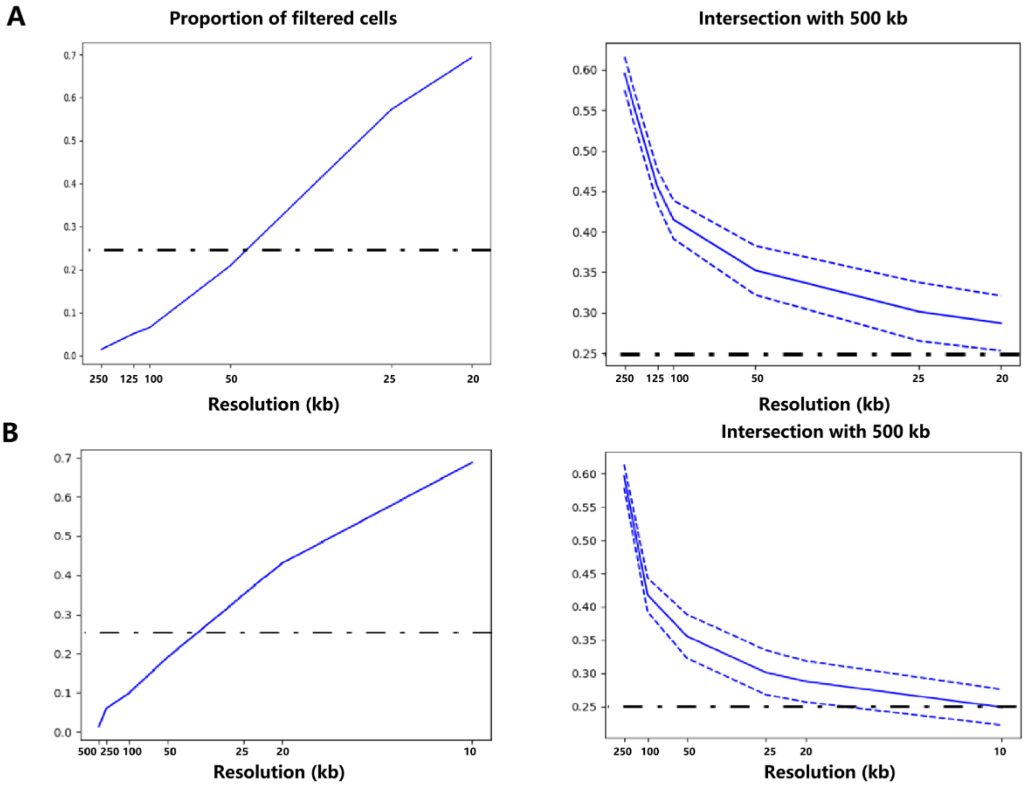


**Supplementary Figure S15. Comparison of subcompartment annotations across different resolutions in the LiMCA and scMicro-C datasets.**

For scMicro-C (A) and LiMCA (B), we plotted the proportion of cells filtered by scSTEC (left) and the intersection between subcompartment annotations at each resolution and those at 500 kb resolution (right). These analyses illustrate the dependence of scSTEC performance on resolution and data quality. Guided by the downsampling analyses in Supplementary Figure S14, we used a filtered-cell fraction below 25% and an intersection above 0.25 as practical empirical criteria. Based on these criteria, the current LiMCA and scMicro-C datasets supported subcompartment annotation at resolutions of approximately 50--100 kb.

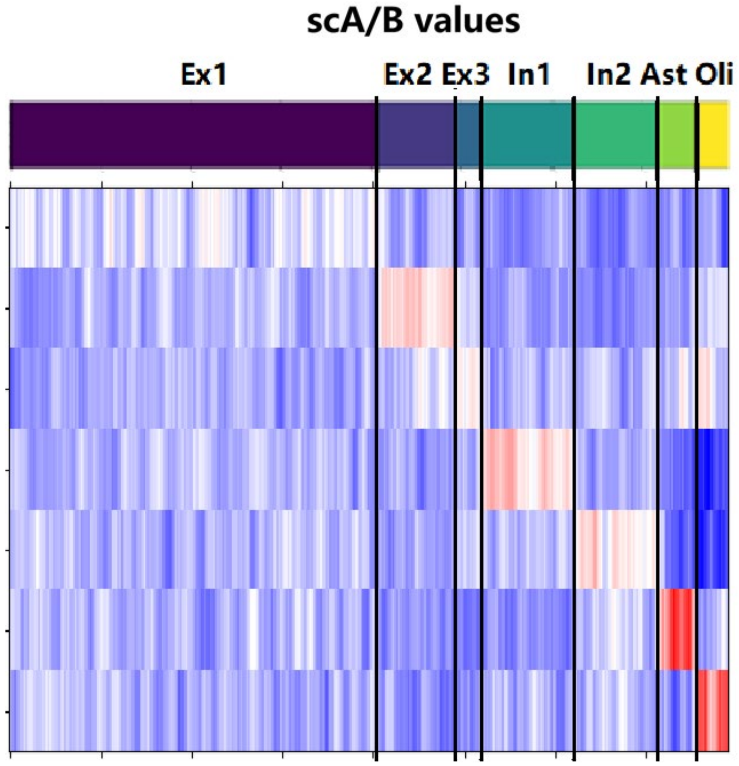


**Supplementary Figure S16. scA/B values at cell-type marker genes.**

We used the top 200 marker genes for each of the seven cell types in the HiRES adult mouse cortex dataset. For each cell, the mean scA/B value was calculated across the marker genes of each cell type. The resulting values were then z-score normalized across cells for each marker-gene set. Each column represents a single cell, and each row represents the marker-gene set of one cell type.

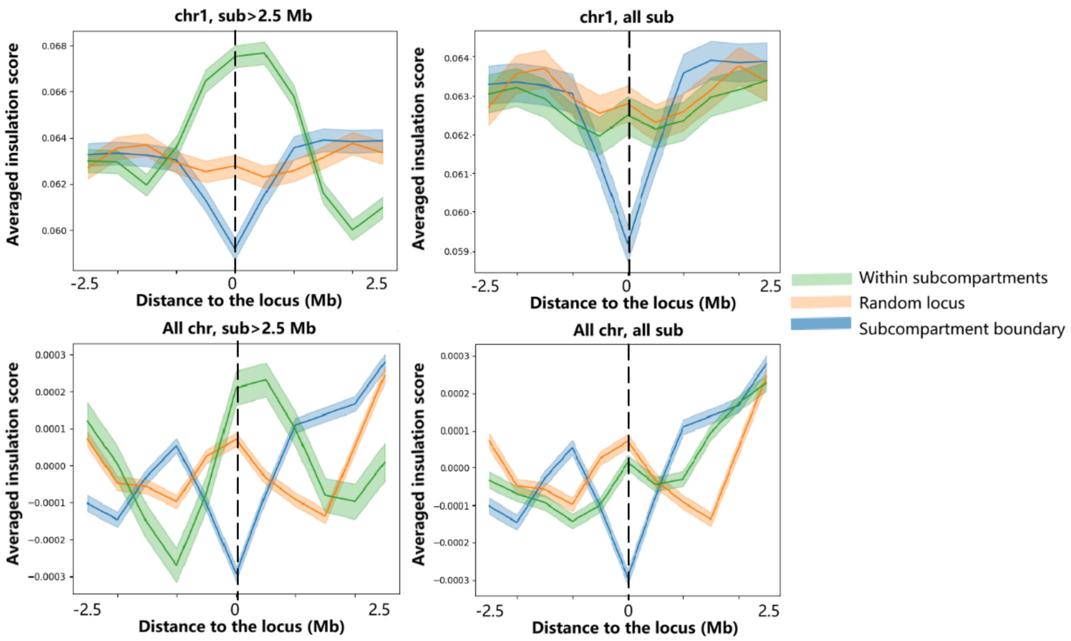


**Supplementary Figure S17. Comparison of subcompartment boundaries with TAD-like domain boundaries in the HiRES adult mouse cortex dataset.**

Aggregated single-cell insulation scores at 500 kb resolution around subcompartment boundaries, subcompartment centers, and randomly selected genomic loci. Insulation scores are shown as mean values with error bars. The top two panels show aggregated insulation scores for chromosome 1, considering subcompartments larger than 2.5 Mb and all subcompartments, respectively. The bottom panels show the corresponding analysis using genomic loci across all chromosomes.

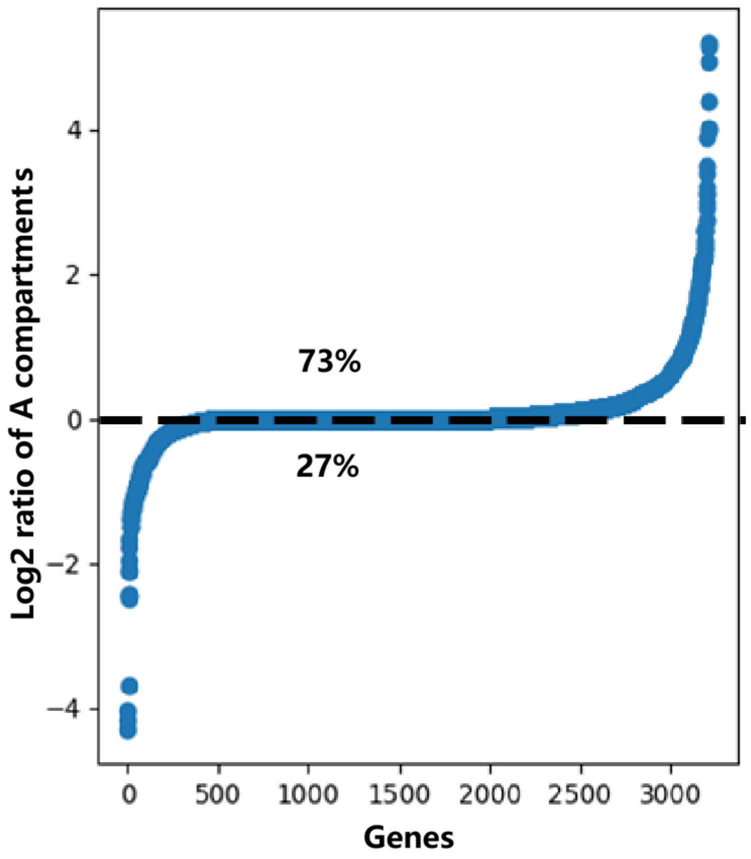


**Supplementary Figure S18. Relationship between single-cell compartment states and transcription within one cell type.**

Using all Ex1 cells in the HiRES adult mouse cortex dataset, we compared the percentage of cells in which the TSS of each gene was assigned to scA (scA1 or scA2) between transcriptionally active cells (UMI>=10) and silent cells (UMI=0). For each gene, the $log_2$ ratio of the scA percentages between transcriptionally active and silent cells was calculated, and genes were ranked according to this ratio.

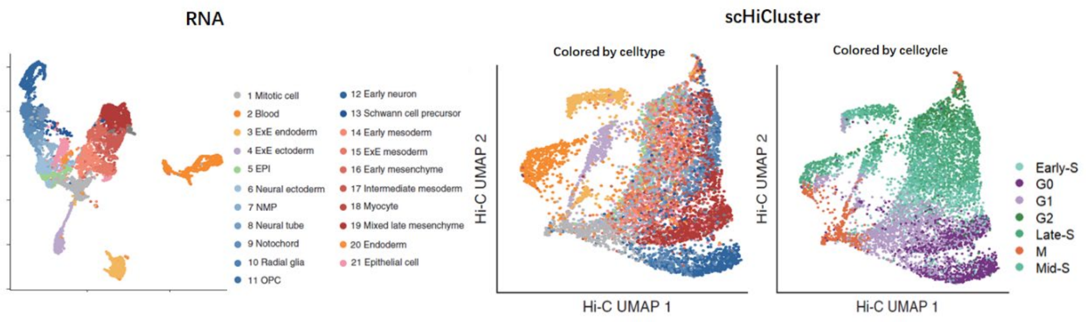


**Supplementary Figure S19. Cellular embeddings based on scRNA-seq and scHi-C data from the HiRES developing mouse embryo dataset.**

Cellular embedding procedures are described in the Supplementary Methods. Transcriptomic embedding was generated from scRNA-seq data, whereas scHiCluster was used to generate the embedding from scHi-C data.

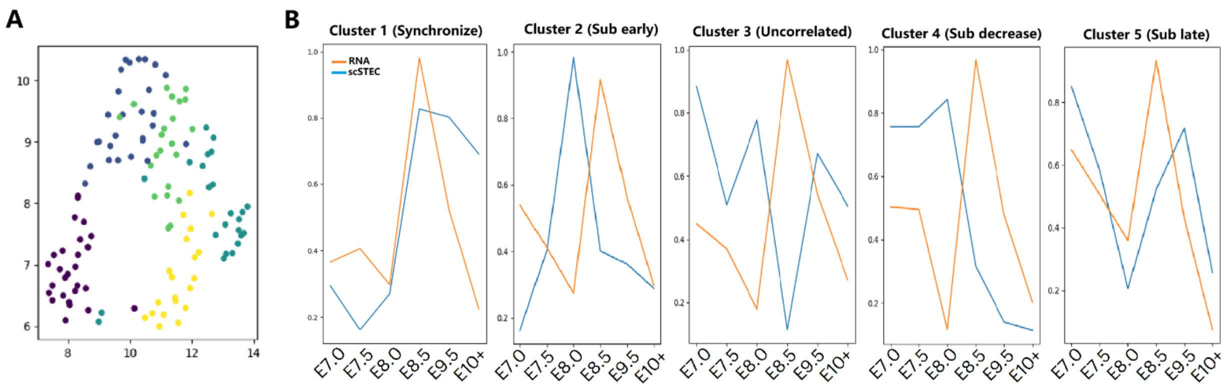


**Supplementary Figure S20. Dynamics of subcompartment states and gene expression along the developmental trajectory for E8.5 marker genes.**

(A) UMAP visualization of E8.5 marker genes based jointly on their subcompartment and gene-expression dynamics along the developmental trajectory. Louvain clustering of the resulting embedding identified five gene clusters.

(B) Trajectory patterns of the five gene clusters identified in (A). The clusters were categorized as Synchronized, Sub early, Uncorrelated, Sub decrease, and Sub late according to the temporal relationships between their subcompartment and gene-expression dynamics.

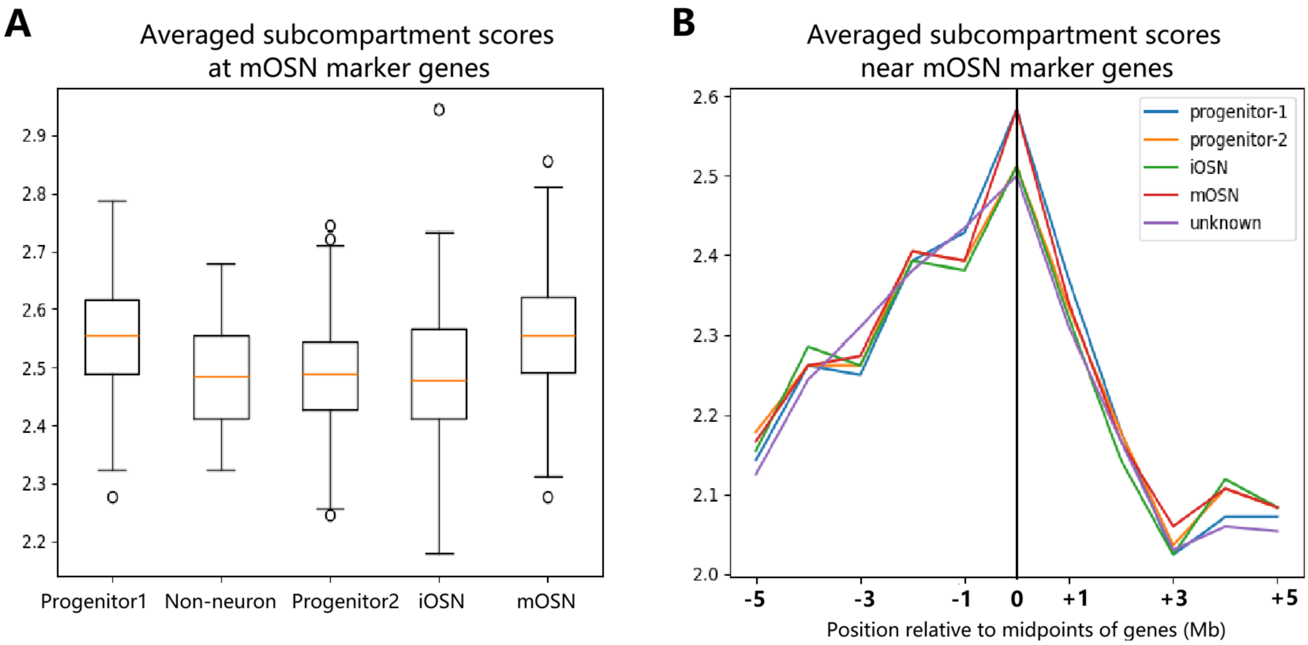


**Supplementary Fig. S21. Subcompartment states of mOSN marker genes and their neighboring regions.**

(A) Mean subcompartment scores across 100 mOSN marker genes reported by Tan et al. (2015) for each cell. Subcompartments were encoded from 0 to 4, corresponding to B3, B2, B1, A2, and A1, respectively.

(B) Subcompartment scores within 5 Mb upstream and downstream of the midpoints of genes (Mb).

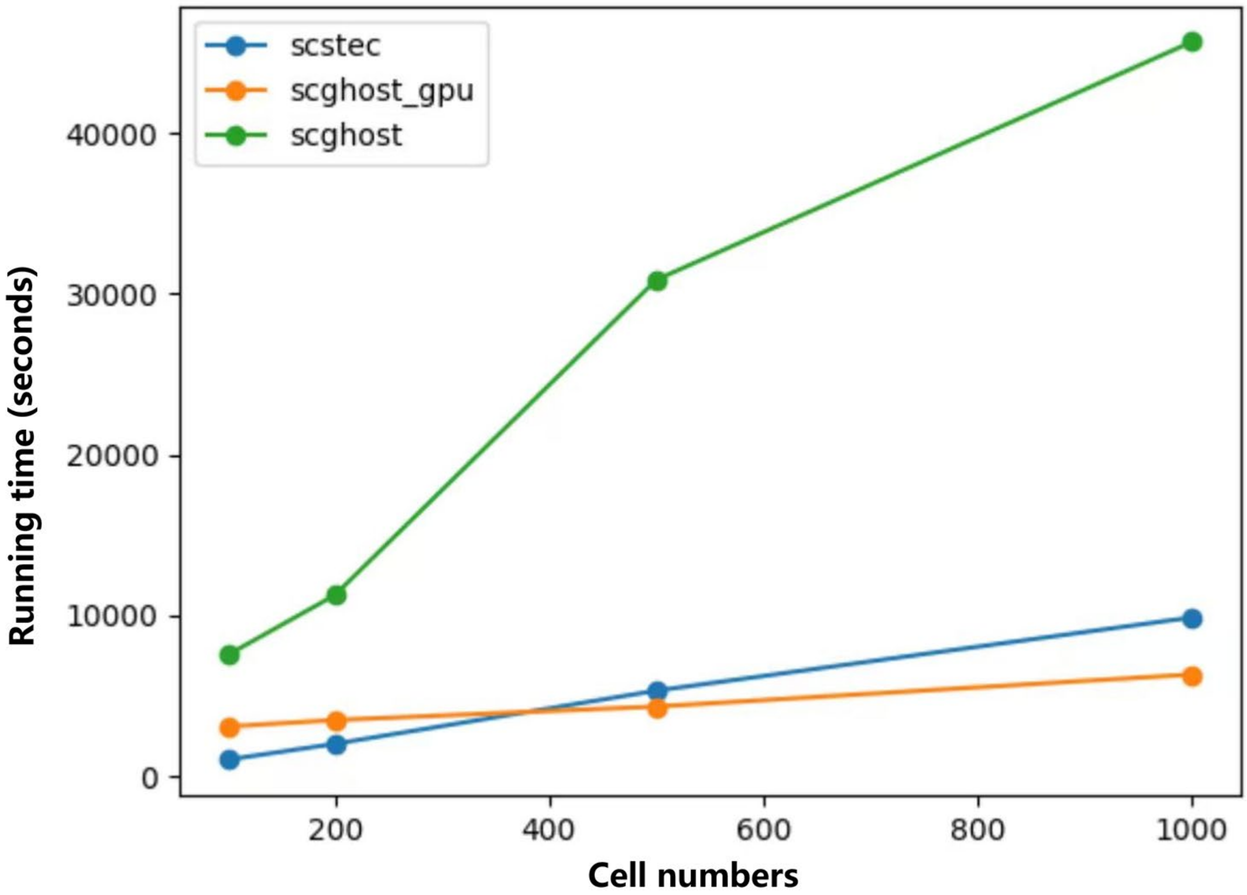


**Supplementary Figure S22. Running time of scSTEC and scGHOST across varying numbers of cells.**

Running time (seconds) of scSTEC and scGHOST with increasing numbers of cells. CPU-based analyses were performed on a machine equipped with Intel Xeon Platinum 8358 processors and 1024 GB of memory using 32 CPU cores. GPU-based scGHOST was evaluated using a single NVIDIA A100 GPU. scSTEC and GPU-based scGHOST required substantially less running time than CPU-based scGHOST, and their running times increased approximately linearly with the number of cells.
